# Integrative analysis of transcriptomics in human craniofacial development reveals novel candidate disease genes

**DOI:** 10.1101/2022.02.28.482338

**Authors:** Tara N. Yankee, Andrea Wilderman, Emma Wentworth Winchester, Jennifer VanOudenhove, Justin Cotney

## Abstract

Craniofacial disorders are among the most common of all congenital defects. A majority of craniofacial development occurs early in pregnancy and to fully understand how craniofacial defects arise, it is essential to observe gene expression during this critical time period. To address this we performed bulk and single-cell RNA-seq on human craniofacial tissue from embryonic development 4 to 8 weeks post conception. This data comprises the most comprehensive profiling of the transcriptome in the early developing human face to date. We identified 239 genes that were specifically expressed in craniofacial tissues relative to dozens of other human tissues and stages. We found that craniofacial specific enhancers are enriched within 400kb of these genes establishing putative regulatory interactions. To further understand how genes are organized in this program we constructed coexpression networks. Strong disease candidates are likely genes that are coexpressed with many other genes, serving as regulatory hubs within these networks. We leveraged large functional genomics databases including GTEx and GnomAD to reveal hub genes that are specifically expressed in craniofacial tissue and genes which are resistant to mutation in the normal healthy population. Our unbiased method revealed dozens of novel disease candidate genes that warrant further study.

## Introduction

Craniofacial disorders are among the most common of all congenital defects, with clefts of the lip and/or palate being the most frequent, affecting nearly 1 in 700 live births worldwide (Leslie and Marazita, 2013). These defects are a significant public health issue with far reaching economic ramifications. For children born with orofacial clefts alone, the combined lifetime cost of treatment is nearly $700 million (Boulet et al., 2009; National Research Council et al., 2012; NIDCR, 2018). Craniofacial disorders also pose unique physiological and psychological challenges to patients and their families. The face contains most of the major sensory organs and is the primary means by which humans communicate emotions, a fundamental social behavior necessary to form and cultivate relationships. For these reasons there is significant impetus to understand the etiology of craniofacial defects to aid in the development of improved diagnostic and preventative methods.

The link between craniofacial abnormalities and genetic disorders has been long established, as over 500 mendelian syndromes (OMIM^®^, 2021) exhibit a craniofacial element. Machine learning technology has been able to take advantage of this observation and several tools have been developed to match facial phenotypes with known syndromes (Gurovich et al., 2019; Huang et al., 2008; Pantel et al., 2020; Taigman et al., 2014). Most recently machine vision tools have shown potential for applications in diagnosing and characterizing ultra-rare syndromes with the added benefit of being minimally invasive (van der Donk et al., 2019; Hsieh et al., 2022). However the majority of craniofacial abnormalities are much more common than these syndromes and occur in the absence of other abnormalities (Leslie and Marazita, 2013; Morris, 2016). The utility of such approaches for diagnosing non-syndromic patients is thus unknown and it remains unclear how many different genes contribute to craniofacial development. Genome-wide association studies (GWAS) have identified dozens of loci that contribute to the heritability of non-syndromic orofacial clefting as well as measures of normal human facial variation. (Adhikari et al., 2016; Beaty et al., 2010; Birnbaum et al., 2009; Bureau et al., 2014; Camargo et al., 2012; Claes et al., 2018; Grant et al., 2009; Leslie et al., 2017; Ludwig et al., 2012, 2016, 2017; Mangold et al., 2010; Mostowska et al., 2018; Yu et al., 2017). A handful of rare variants have been identified in genes, however most are found in non-coding portions of the genome. We and others have shown that such variants are systematically enriched in sequences that obtain active chromatin states during early human craniofacial development, particularly those annotated as strong enhancers (Bonfante et al., 2021; Liu et al., 2021; Wilderman et al., 2018). Strong enhancers only active in developing craniofacial tissues were enriched for craniofacial related transcription factor (TF) binding sites and generally localized near genes implicated in craniofacial abnormalities. However these enrichments were driven by a relatively small fraction of all identified craniofacial enhancers and establishing enhancer-gene assignments is difficult in the absence of direct gene expression measurements (Fulco et al., 2019; Moore et al. 2020; Whalen et al., 2016).

Thus far gene expression during craniofacial development has been primarily studied in model organisms such as mouse, chicken, and zebrafish (Askary et al., 2017; Brugmann et al., 2010; Chang et al., 2014; ENCODE Project Consortium, 2012; Fabian et al., 2022; Hooper et al., 2017; Li et al., 2019; Potter and Potter, 2015). Due to ethical and logistical reasons, gene expression data from primary craniofacial developing human tissue are rare (Cai et al., 2005; Samuels et al., 2020) compared to the thousands of gene expression data sets produced in adult tissues by large-scale consortia such as Encode and GTEx. Thus there is great need for large-scale genome-wide data in order to comprehensively study human craniofacial development and better understand which genes are regulated by the trait loci identified in patient cohorts. To address this knowledge gap we have generated an extensive set of bulk RNA-seq data from organogenesis of the human face.

Here we describe analysis of these gene expression data along with integration of thousands of gene expression profiles from adult tissues and our previously published craniofacial chromatin state data. These analyses revealed genes with previously unappreciated craniofacial-specific expression and allowed us to predict target genes of craniofacial-specific enhancers. We show how multiple layers of genomic data can be used to provide a biological frame of reference for SNPs identified by GWAS or Whole Genome Sequencing (WGS). We also use this analysis to predict a set of novel genes specifically important for craniofacial development. We further characterize several of these genes to reveal differences in cell-type specificity between human and mouse. These resources are available from the Genome Expression Omnibus, dbGAP, as a public track hub on the University of California Santa Cruz (UCSC) Genome Browser, and directly from our laboratory website (https://cotney.research.uchc.edu/craniofacial).

## Results

### Characterization of gene expression during human craniofacial development

Gene expression during human organogenesis has been shown to be highly dynamic and its precise orchestration is crucial for normal patterning of a variety of tissues and structures (Cardoso-Moreira et al., 2019; Mazin et al., 2021; VanOudenhove et al., 2020). However, gene expression patterns during critical stages of human craniofacial development have not been previously profiled. To fully characterize the transcriptome of the developing human face, we generated bulk RNA-seq libraries on primary craniofacial (CF) tissue from four distinct Carnegie Stages (CS) of the embryonic period (CS13, CS14, CS15, CS17) and a culture model of cranial neural crest cells (CNCCs). We also retrieved CS22 data (Pennacchio et al., 2017) from Facebase resulting in a time series data set encompassing nearly half of the embryonic period, including most of the critical events in CF development (Figure 1A). Quality control metrics revealed newly generated samples had minimal 3’ bias, generally high RNA integrity number (RIN) values (>8), and excellent transcript integrity number (TIN) scores (>70) (Figure 1B; S1A, Table S1). Principal component analysis (PCA) of the novel CF samples showed minimal effects from sex and RNA quality in the first 2 components (Figure S1B). Having confirmed the high quality of these samples, we then summarized gene expression at the composite gene level and identified an average 26000 genes expressed at each timepoint (see Methods). The greatest number of genes were expressed at the latest timepoints of the developmental series, consistent with the increasing heterogeneity of tissues and cell types as development progresses. This trend was further demonstrated by comparison of global gene expression profiles, which revealed the first major principal component of gene expression coincided with developmental time (Figure 1B).

**Figure 1.**
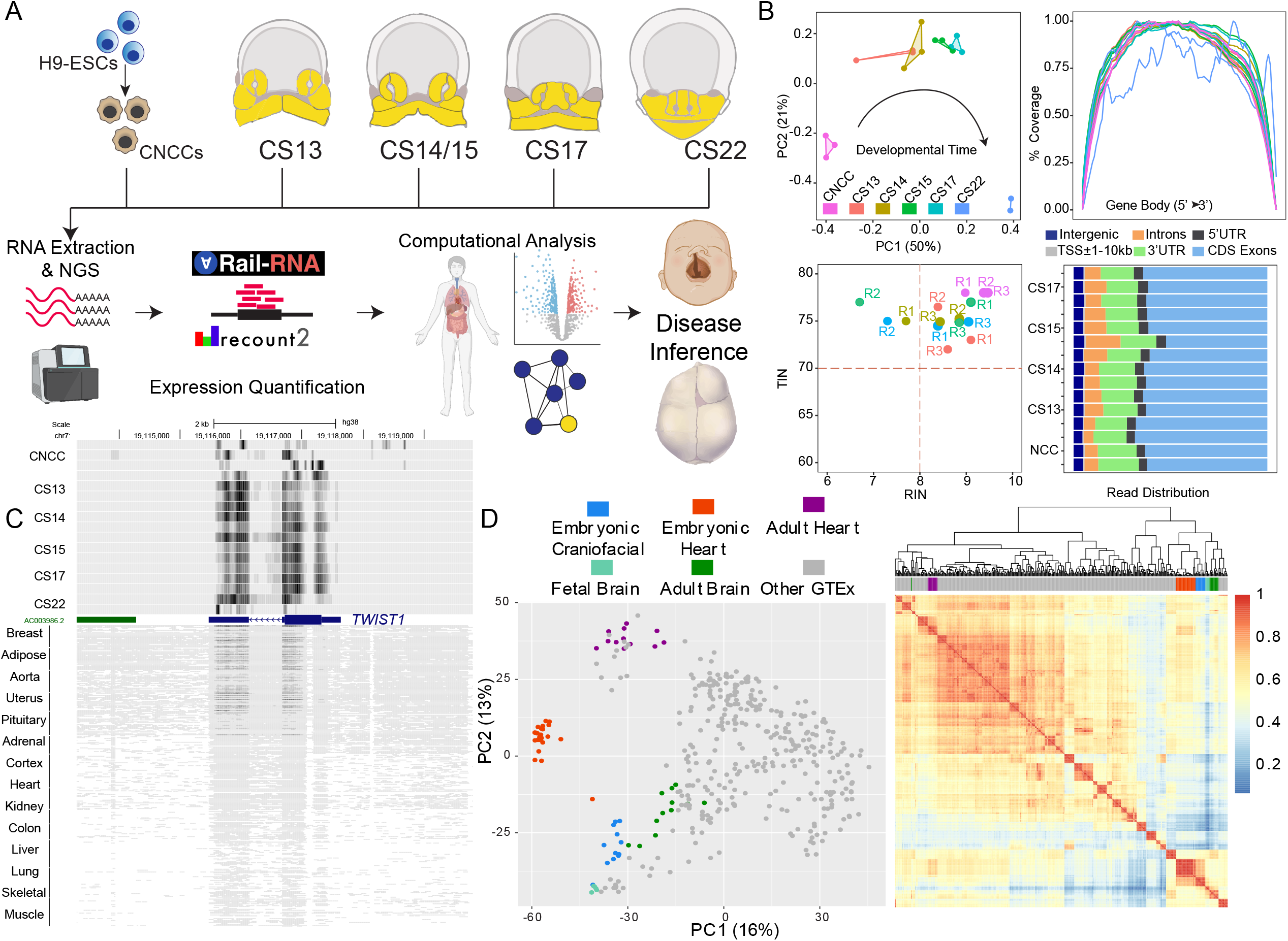
Transcriptomic Profiling of Early Human Craniofacial Development. (A) Overview of experimental design. Bulk mRNA-seq was performed on five time points of the primary CF tissue and a cell culture model of CNCCs. The Rail-RNA/recount2 pipeline was used for alignment and gene-level transcript quantification. In order to determine potential disease genes within craniofacial development several downstream computational analyses were performed such as multi-tissue comparisons, pairwise differential expression and the generation of co-expression networks. (B) Quality control metrics. *Upper left* Gene body coverage plots show reads lack a 3’ bias. *Upper Right* Read distribution plots show reads are dominated by CDS (coding) Exons. *Lower left* RIN vs. TIN plot. TIN values established by (Wang et al., 2016) were calculated using RSeQC and are all above the recommended threshold of 70. *Lower right* The PCA plot shows that the samples vary the most by time which is captured by PC1. (C) Genome browser shot of *TWIST1* bulk RNA-seq signal across several tissues. TWIST1 expression is heavily biased in the craniofacial tissues (Figure S1E). (D) Multi-tissue comparison between the early primary craniofacial tissue, embryonic heart, fetal brain and 12 random samples from each of the primary tissues available in GTEx. *Left* PCA plot of the top 500 most variable genes. *Right* Spearman correlation of global gene expression.

While these initial comparisons gave us a sense of the overall trends in the data, it was unclear if there might be systematic biases in our data due to the source of the material and methods of preservation. To address this issue we uniformly processed our data with the *Rail-RNA* pipeline (Nellore et al., 2017) employed by the *recount2* database (Collado-Torres et al., 2017) to enable direct comparisons of gene expression across thousands of samples collected from tissues profiled by GTEx and the developing human heart and brain (Fietz et al., 2012; Melé et al., 2015; VanOudenhove et al., 2020). When we compared composite gene level expression across all of these samples we observed similar numbers of expressed genes and ranges of gene expression values across all samples (Figure S1C). When we inspected the developmental transcription factor *TWIST1*, implicated in autosomal dominant craniosynostosis Saethre-Chotzen syndrome (Cai et al., 2003), we observed robust expression in CF samples and relatively lower levels in all other human tissues (Figure 1C, S1E). We then performed a tSNE analysis on all expressed genes across all samples and found a distinct cluster of CF samples, along with clusters for each of the tissues profiled by GTEx (Figure S1D). This trend was also observed in PCA, which showed CF tissues generally clustered together. Quantitatively, pairwise correlations between samples revealed a global delineation in prenatal and postnatal gene expression (Figure 1D). Overall, these results demonstrate the quality of our data, the robustness of individual embryo staging and recovery of a clear developmental time component in gene expression.

### Genes expressed specifically during craniofacial development are enriched for craniofacial abnormalities

Specificity of gene expression is an important aspect of cell identity in multicellular organisms. Restricted gene expression is frequently observed for genes directly related to tissue-specific functionality and is often an indicator for disease relevance (GTEx Consortium, 2017; Kitsak et al., 2016; Lage et al., 2008). Our previous work has shown that specificity of expression directly identified known genes for congenital heart defects (VanOudenhove et al., 2020) and as demonstrated above, *TWIST1* seemed to be specifically expressed in our CF samples.

To systematically identify genes with similar patterns of tissue-specific expression, we used a measurement of inequality called the Gini Index across all tissues profiled above. (Ceriani and Verme, 2012; Gini, 1912, 1914) (Table S2). We identified over 5,000 “Gini” genes that have highly restricted expression (Gini Index >=0.7) (Figure 2A). Of these, 239 genes demonstrated the highest average expression in our CF samples. To determine the disease relevance of these genes, we performed a disease enrichment analysis of these CF Gini genes and found they were most enriched for CF abnormalities and cleft palate. Consistent with these findings of Gini gene tissue relevance, fetal brain and embryonic heart Gini genes were significantly enriched for neurological disease and congenital heart defect related ontologies respectively (Figure 2B).

**Figure 2.**
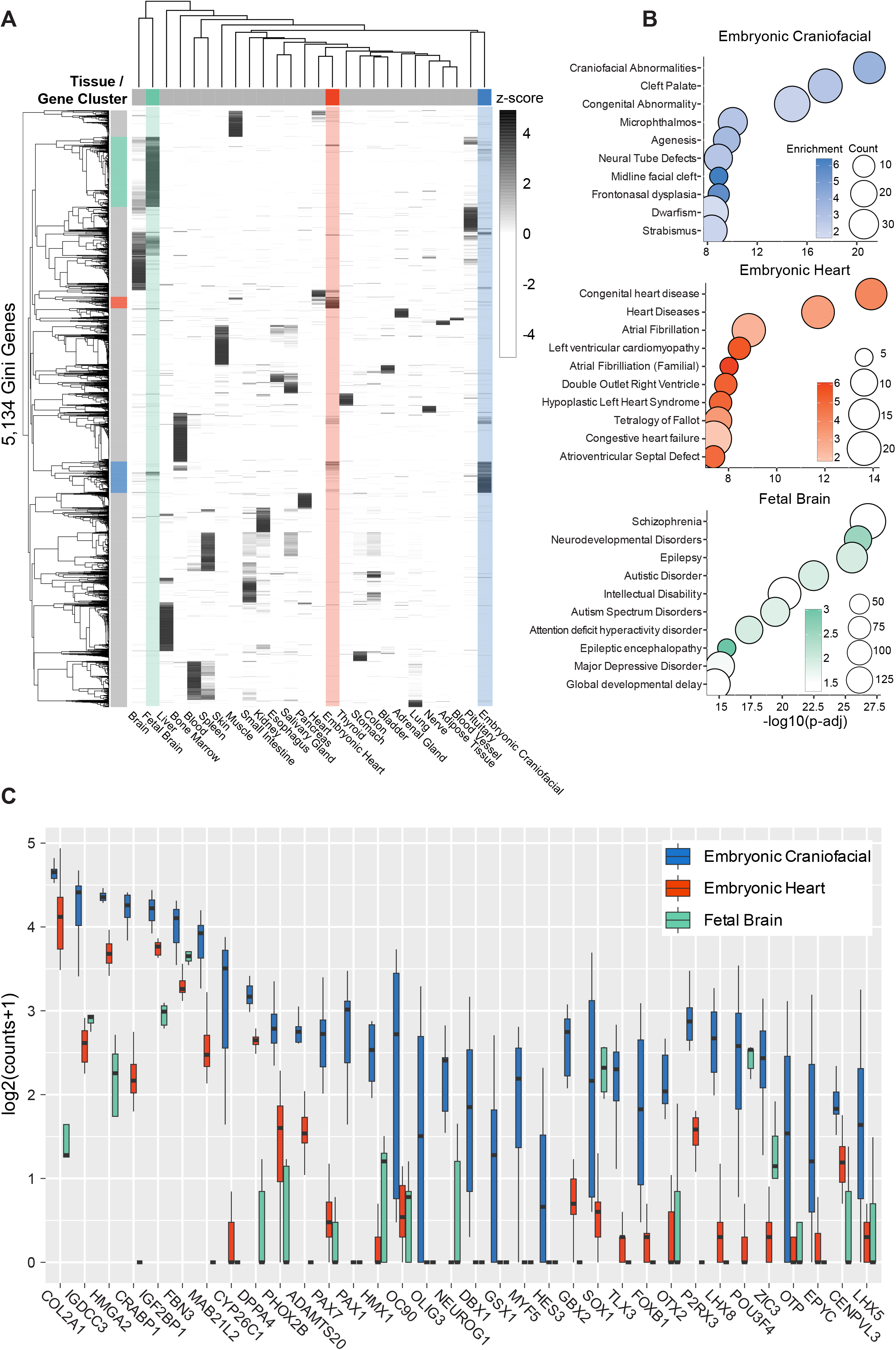
Gini Index unbiasedly identifies gene expression specific to embryonic craniofacial tissue genome-wide. (A) Heatmap showing specificity of expression for tissue specific genes (rows) identified with high Gini scores (>=0.7) for all non-sex (n=25) tissues (columns) from GTEx, embryonic heart and fetal brain. Gene expression increases from light to dark. Embryonic craniofacial (blue), embryonic heart (red) and fetal brain (light green) specific genes and samples are highlighted, all other tissues are marked gray. (B) Selected significant (Benjamini and Hochberg–adjusted p value <0.05) disease ontology enrichments for genes identified to be specific for these tissues (Gini >=0.7 and highest average expression). The dot size or Count indicates number of genes within a disease category. Color bars are log2 of the fold enrichment (see Methods). (C) Boxplots of gene expression of highest scoring Gini genes for the craniofacial tissues compared to embryonic heart and fetal brain.

Many transcription factors (TFs) have restricted patterns of expression and are critical in developmental patterning and specification across many tissues (Lambert et al., 2018). Therefore we asked if CF Gini genes are enriched for TFs. Amongst the CF Gini genes we identified a significant enrichment of TFs (n=71, Fisher exact p< 2.2X10^-16) including many known to be important in CF development such as *ALX1, ALX4, DLX5, DLX6, PAX3* and *TWIST1 (Gou et al., 2015)*. In addition to transcription factors, signaling pathways are also well appreciated to be important for normal CF development. Consistent with this established biology, we found significant enrichment of genes belonging to families of signaling molecules known to be important in regulating CF morphogenesis including *FGF8, FGF19* (Kurose et al., 2004; Wang et al., 2013; Weng et al., 2018) and *WNT7A, WNT7B, & WNT8B* (Figure S2A) (Reynolds et al., 2019).

The genes with the highest Gini index for CF tissues are shown in Fig 2C. The most highly expressed, top scoring gene *COL2A1* has been linked to Stickler syndrome which includes CF abnormalities like cleft palate and midline facial hypoplasia (Hoornaert et al., 2010; Kondo et al., 2016)). While this approach successfully revealed genes known to drive CF disease, the unbiased nature of this analysis also allowed us to uncover candidate genes not previously associated with CF development. For instance *HMGA2* is one of the most specifically expressed genes in our CF tissues. This gene has been linked in OMIM to Silver-Russell Syndrome patients and reported in a pygmy mouse phenotype, but is relatively underappreciated in CF development (Federico et al., 2014; Zhou et al., 1995). Another top scoring gene in our analysis, *IGF2BP1*, encodes a RNA-binding protein (RBP) that has not been linked to any human diseases in OMIM but results in specific cartilage loss in nasal bones and mandible in mouse (Hansen et al., 2004). RBPs as a class have been linked to a handful of syndromes with CF phenotypes, suggesting *IGF2BP1* may also play a key role in CF development in humans (Lehalle et al., 2015). Overall, Gini analysis captured many known CF disease genes and provided many novel genes that warrant further inspection in human CF development. Moreover, this analysis identified thousands of genes with restricted expression across tissue types which are likely important to many different fields.

### Integration of chromatin states and gene expression from multiple tissues reveal trends in regulatory architecture for tissue specific genes

Given the enrichment of genes with known roles in craniofacial development and significant numbers of transcription factors in our Gini analysis, we asked if tissue-specific regulatory sequences might play a role in these restricted expression patterns. Non-coding regulatory regions such as enhancers are known to act on genes in a spatiotemporal manner (Panigrahi and O’Malley, 2021) and accessible regulatory regions have been shown to be enriched for disease-associated variants (Maurano et al., 2012; Thurman et al., 2012). In previous work we have shown that enhancers active in human craniofacial and heart development are enriched for variants associated with risk for orofacial clefting and atrial fibrillation respectively, suggesting they act on tissue-specific developmental genes (VanOudenhove et al., 2020; Wilderman et al., 2018).

To determine if CF Gini genes are likely regulated by CF-specific enhancers we leveraged epigenetic atlases from embryonic development along with fetal and adult tissues in Roadmap Epigenome (Roadmap Epigenomics Consortium et al., 2015). Using the “strong” enhancer states identified from the 25 ChromHMM analysis (Ernst and Kellis, 2017), we established a list of 10,693 putative novel CF-specific strong enhancer segments (CFSEs) by comparing to all other available tissues in Roadmap Epigenome and embryonic heart (Table S3). Using GREAT to identify CFSE gene targets, we found that as the number of CFSEs targeting a gene increased, the level of gene expression increased specifically in CF tissue compared to other embryonic, fetal, and adult tissues (Figure 3A). These results suggest that CFSEs drive tissue-specific gene expression and many are likely to be directly targeting these genes *in vivo*. We indeed found CF Gini genes had the largest enrichment of CFSE target genes, followed by fetal brain and embryonic heart confirming the global gene expression trends between tissues (Figure 1D).

**Figure 3.**
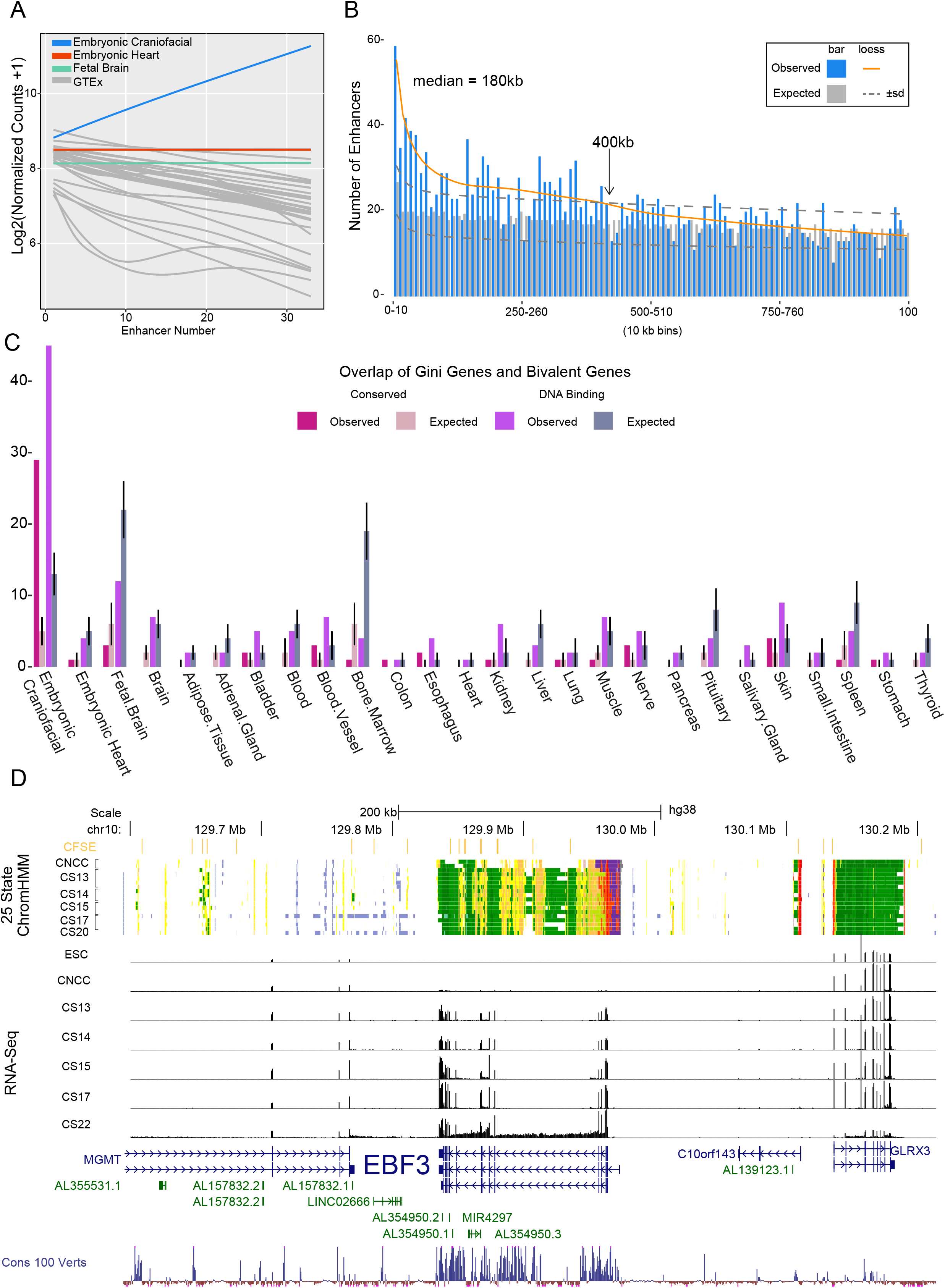
Strong novel craniofacial enhancers target Gini genes. (A) Expression values versus CFSEs number modeled by linear loess smoothing for embryonic craniofacial (blue), embryonic heart (red), fetal brain (green) and all tissues in GTEx (gray). (B) Histogram plot of distance for closest craniofacial (blue bars) Gini gene to CFSEs compared to a randomly picked Gini gene from all other tissues (gray bars). The median and standard deviation were calculated from 1000 iterations. Each line is an interpolation for craniofacial Gini genes (orange) and for the maximum and minimum random Gini genes (dark gray dashed). (C) Gini gene enrichment for genes with bivalent promoters in craniofacial tissue. The pink bars are bivalent genes conserved between human and mouse, and the purple bars are bivalent regions spanning DNA-binding genes. The median and standard deviation were calculated from 1000 iterations of randomly selected genes. (D) Genome browser shot of *EBF3* locus showing 25 state ChromHMM segmentation from culture model of CNCCs, and CS13 through CS20 of primary human craniofacial tissue. EBF3 has a bivalent promoter in these samples (dark purple). Above the segmentation tracks is the active and novel CFSE track shown in orange which include strong enhancer ChromHMM states 13-15. Below are RNA-seq bigwig signal tracks for human embryonic stem cells (ESC), CNCCs, and CS13 through CS22.

Given this robust association between CFSEs and CF Gini genes, we asked where tissue-specific enhancers tend to be physically located relative to their targets. Enhancers have been demonstrated to form long-distance interactions with their target genes, with some of the most relevant and specific interactions occurring over distances greater than 1 Mb in developing tissues (Lettice et al., 2002, 2003). Conversely in cultured cells, proximity between high confidence gene-enhancer pairs was strongly enriched within 50 kilobases of a TSS (Gasperini et al., 2019). These contrasting findings raised the possibility that very long range enhancer-gene interactions are more common in development. To address this question we assigned each CFSE to the closest CF Gini gene *in cis*. Compared to a random permutation control, we detected significant enrichment of CFSE-Gini pairs at distances up to 400kb with a median distance of 180kb. Approximately 20% of these CFSEs were located within 50 kb of a CF Gini gene and in total were significantly further away than high confidence enhancer-gene pairs identified via CRISPRi in K562 (Figure 3B, S3A-B). These results suggest enhancer targeting events in development are enriched for longer distances, which is consistent with previous observations that developmental TFs are often flanked by large non-coding regions (Laverré et al., 2022; Ovcharenko et al., 2005) resulting in longer distances for enhancer targeting events. However extremely long range interactions (>500kb) seem to be relatively rare and do not rise above background expectation.

The results above indicated tissue-specific, active chromatin states play strong roles in specificity of expression. However, repressive chromatin modifications and the complexes that deposit those marks have been associated with developmental abnormalities. Specifically, components of the Polycomb Repressive Complex 2 (PRC2), particularly EZH2 that catalyzes H3K27me3 deposition, are essential for normal mouse development and have been implicated in neural crest derived tissue specification as a result of global gene derepression (Boyer et al., 2006; Bracken et al., 2006; O’Carroll et al., 2001; Pasini et al., 2004; Tien et al., 2015). The H3K27me3 mark is strongly associated with bivalent chromatin state segmentations across multiple tissues. Bivalent domains are thought to be regions within the genome that facilitate temporal expediency during development but also identify genes with restricted or gradients of expression within a tissue (Blanco et al., 2020; Roadmap Epigenomics Consortium et al., 2015). Our previous work identified 957 genes with bivalent promoters exclusive to embryonic craniofacial tissue (Wilderman et al., 2018). Many of these were DNA binding factors that are known to have restricted patterns of gene expression across the mouse embryo, including the adjacent transcription factors *DLX5* and *DLX6*, suggesting a relationship between bivalent status and specificity of expression (Acampora et al., 1999; Depew et al., 1999; Robledo et al., 2002). When we examined all CF specific bivalent genes we did not find a correlation with tissue-specificity of gene expression (Figure S3C). However, bivalent genes classified as DNA binding factors were significantly enriched for embryonic CF Gini genes. This trend was also observed for all bivalent promoters conserved between mouse and human developing CF tissues (Figure 3C).

Notably we found that DNA-binding factors with bivalent promoter status during human CF development showed the strongest trends in both the absolute number of assigned CFSEs and the greatest overall specificity of expression. For example the nuclear receptor *NR2F1* gene, which has been shown to be a key regulator of neural crest specification (Rada-Iglesias et al., 2012), is potentially targeted by 16 CFSEs. Other well known CF TFs like *MSX2* and *ALX4* are targeted by 10 or more CFSEs (Jabs et al., 1993; Kayserili et al., 2009; Mavrogiannis et al., 2006; Satokata et al., 2000)(Figure S4).

This analysis also identified several genes not reported to play a direct role in human CF development based on OMIM annotations including *BCL11A, EBF3*, and *HMGA2* (Table S3, Figure S5). In the case of *EBF3* there are 19 CFSEs within 400kb of the bivalent TSS region and we observed CF-specific patterns of expression compared to surrounding genes (Figure 3D). Overall, these results demonstrate a clear relationship with the genomic organization of tissue-specific enhancers relative to genes that have high specificity of expression during development, particularly those that maintain a bivalent state during organogenesis.

### Differential expression during craniofacial development identifies temporal-specific biological processes

Our analysis thus far has demonstrated the value of comparing across multiple tissues for observing global trends of gene expression and identification of genes with tissue-specific patterns of expression. However these analyses assume each tissue is at one time point and thus do not capture gene dynamics throughout development. Craniofacial development is a morphologically complex process which is likely to be reflected in its gene expression, thus we sought to understand how gene expression patterns change throughout early development in craniofacial tissues. We first performed differential expression analysis in a pairwise fashion across all polyA captured RNA from primary tissue time points. We found a total of 3677 genes significantly differentially expressed. The largest number of differentially expressed genes (DEGs) were found in comparisons between the earliest and latest stages (CS13 & CS17, n =2733) consistent with progression of morphogenesis of a structure into multiple distinct tissue types (Figure S6A-C, Table S4).

When we compared the expression of these DEGs across all of our gene expression data we found that hierarchical clustering separated samples into two main groups that we denote as Early (CNCC, CS13 and CS14) and Late (CS15, CS17 and CS22) (Figure 4A). This grouping of samples indicated the culture model of CNCC utilized in this study does indeed reflect the primary tissue when neural crest cells are thought to be populating the craniofacial structures. This is in contrast to an alternative cell culture model which does not share much gene expression dynamics with the primary tissue (Figure S6D).

**Figure 4.**
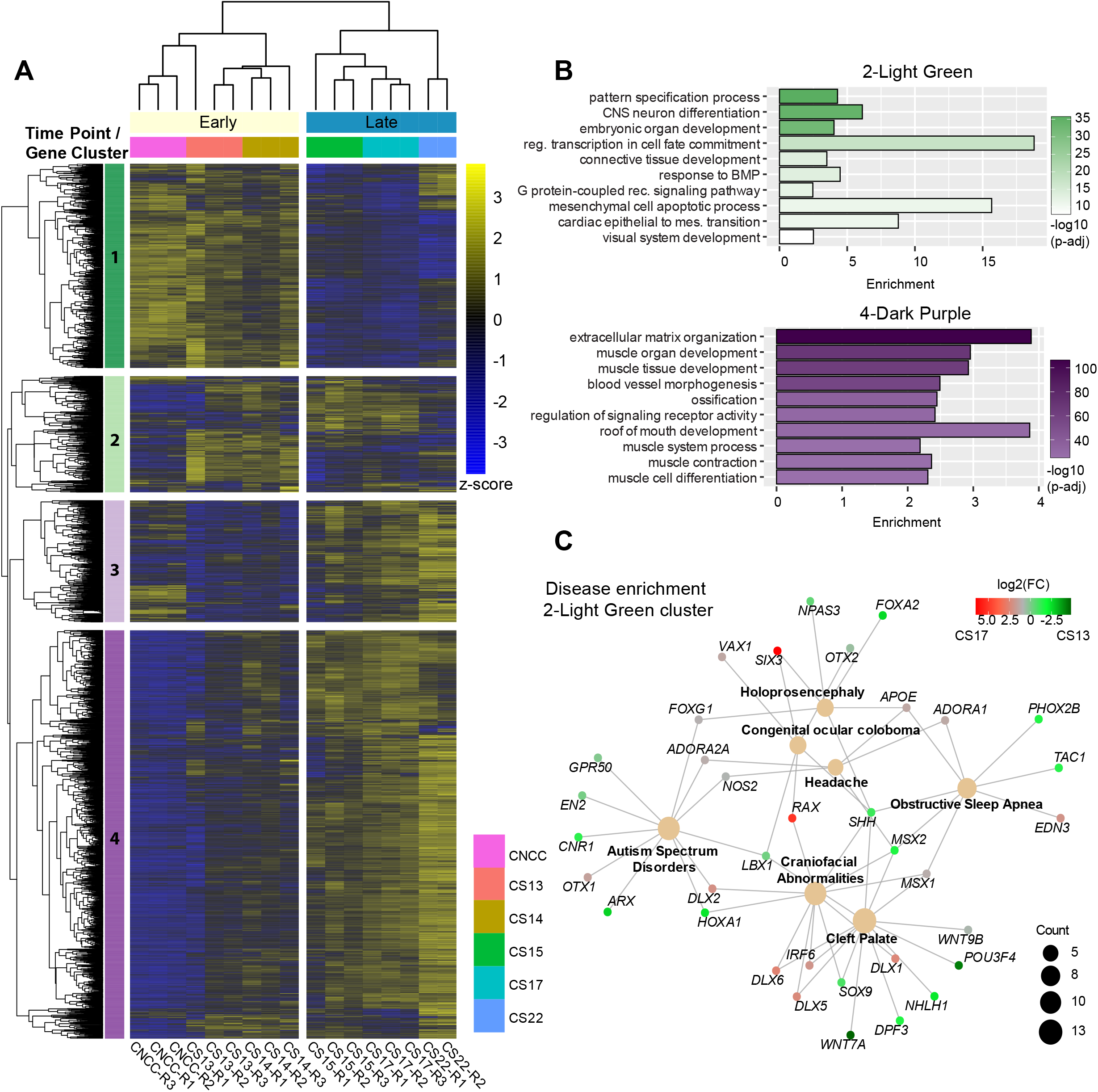
Differential expression during craniofacial development. (A) Heatmap of differentially expressed genes (log_2_ Fold Change > abs(1), Benjamini-Hochberg p-adj <0.05) across all samples in our developmental series. Z-scores for normalized gene counts ranging from high (yellow) to low (blue) expression. Dendrogram on the top is hierarchical clustering of all samples. Samples separated into two main clusters, Early (CNCC,CS13,CS14) and Late (CS15,CS17,CS22). Dendrogram on the left is hierarchical clustering of gene counts. The genes were assigned groups identified by a color and number (1-Dark Green, 2-Light Green, 3-Light Purple, 4-Dark Purple) by cutting the dendrogram at a height which would result in 4 clusters. The genes in the green clusters are for those primarily expressed in the early and intermediate stages of the series. While the purple cluster genes tend to be active in the late stages. (B) Gene ontology of a select subset of the most significant terms identified by clusterProfiler for 2-Light Green and 4-Dark Purple cluster. The genes of the 2-Light Green cluster enrich for embryonic patterning and terms concerning the mesenchyme. The 4-Dark Purple cluster includes muscle developmental functions with ossification and blood vessel morphogenesis. (C) Network plot of selected significant disease ontology terms from genes of the 2-Light Green cluster. Count refers to number of genes within a category and the colorbar shows log_2_ fold change as calculated by DESeq2 between CS13 and CS17 samples which color the gene nodes.

We identified four major clusters of genes which had obvious expression trends during the developmental trajectory (Table S4). The upper cluster (1-Dark Green) showed strong expression in CNCC and CS13 samples and progressive dampening of expression over developmental time. The genes in the upper mid cluster (2-Light Green) are largely absent in CNCC and CS22 but have dynamic expression between early and late stages. These two clusters were enriched for genes involved in patterning, morphogenesis, and cell fate specification (Figure 4B, S6E). Disease enrichment analysis of the light green cluster revealed strong enrichment for CF abnormalities including cleft lip with or without cleft palate (Figure 4C). Genes in the lower mid cluster (3-Light Purple) were expressed in CNCC but largely absent from the rest of the early tissue samples. They are expressed again at CS15 and generally increase across the rest of the developmental trajectory (Figure 4A, S6E). Genes in the bottom cluster (4-Dark Purple) are mostly not expressed in the early stages but progressively increase in expression over the duration of the developmental trajectory. These two clusters were enriched for GO terms involved with cell-cell adhesion, muscle development, ossification, and upper palate development (Figure 4B). These DEGs included all members of the RUNX family of TFs, many members of the myogenic TF family, and all members of the PITX family of TFs.

Several genes identified to be involved in establishing neural crest identity (Simões-Costa and Bronner, 2015) were found in all but one of the clusters, with most residing in dark purple. Interestingly, *PAX3, PAX7, RXRG*, and *SOX9* exhibit oscillating trajectories, which varied from the overall trajectory among all TFs (Figure S7A-B). This may be due to multiple waves of neural crest populating the human CF tissue as it is thought to happen in trunk sensory neurogenesis (Bhatt et al., 2013). These data suggest that the embryonic period of development, particularly the 4 to 8 post conception week period in humans, has distinct programs that are employed sequentially as morphogenesis and organogenesis of the face and skull progress.

The classes of genes linked to disorders with CF abnormalities are exceptionally varied. They include TFs, cell-cell signaling, RNA-binding and protein metabolism (Twigg and Wilkie, 2015; Weinberg et al., 2018). We asked if certain classes of genes would be biased toward dynamic expression in our CF tissues. We found that signaling and TF classes were more prominent in our differentially expressed genes than expected (Figure S7C) while RBPs were depleted. Many RBPs are known to be ubiquitously expressed and we show that they have generally low Gini scores in our multi-tissue analysis (Figure S7D). It is therefore likely that their function is not directly regulated by transcription, but instead may be important for post-transcriptional processing of genes that are either specifically expressed or expressed in cell types like migrating neural crest that are particularly sensitive to perturbation (Forman et al., 2021).

Based on our above multi-tissue analysis, specifically expressed genes and TFs with bivalent chromatin status are indicated to play important roles in CF development. We found that CF Gini genes were most enriched in the early green clusters (Fisher exact p< 2.2X10^-16). Bivalent transcription factors showed similar trends in enrichment across clusters with the most significant enrichment in the light green and dark purple clusters (Fisher exact p<2.2X10^-16). Suggesting that throughout development there are several narrow windows of time when craniofacial specific biological processes are occurring.

When we analyzed CS13 chromatin states at the promoters of genes in each cluster we found consistent trends in activating histone modifications, particularly H3K4me3 levels of the dark green cluster that are more strongly expressed in the early time period (Figure S8). Additionally when we inspected differential enhancer utilization across development we found enhancers of DEGs showed similar trajectories of H3K27ac levels (Figure S9). Overall the patterns of expression that were revealed in the differential expression analysis were largely confirmed by chromatin state both at promoters and distal enhancers suggesting direct regulation of genes by changes in histone modifications at these classes of regulatory sequences.

### Coexpression networks reveal groups of genes with distinct roles in craniofacial development

The differential expression analyses above revealed clear dynamics in gene expression across a subset of genes. However genes not rising to stringent pairwise differential expression criteria are not considered, which potentially misses information about other important gene relationship patterns such as coexpression. Gene coexpression during development has been shown to reveal modules of genes that have similar patterns of regulation across time and even identify cell-type specific enrichments from bulk data (Chatterjee et al., 2019; VanOudenhove et al., 2020; Werling et al., 2020). Coexpression network analysis has been successfully applied in the developing brain, particularly for unraveling the complex genetic architecture of Autism Spectrum Disorder (ASD) (Cotney et al., 2015; Parikshak et al., 2013).

Given that craniofacial disorders are congenital, complex and common we wondered if the same trends might exist in our expression data and reveal novel disease gene candidates. To uncover any such genes, we generated coexpression networks using weighted gene coexpression network analysis (WGCNA) which has been employed in both human brain and heart development (Langfelder and Horvath, 2008). After normalization and filtering for low expression, 26626 genes were assigned to 29 modules ranging in sizes from ~150 to over 4,000 coexpressed genes. We hypothesized that there would be modules unique or important to CF development, and to help identify them we performed GO enrichment (Figure 5A). The GO analysis revealed that the modules were biologically distinct from one another (Table S5). The decreasing modules captured by WGCNA were enriched for many generic cellular processes such as cell cycling (orange), tRNA processing (green), splicing (darkgreen) and protein trafficking (darkorange). Amongst the increasing modules more tissue-specific enrichments were observed including muscle development (darkmagenta), epithelial differentiation (lightyellow), and most notably embryonic patterning and morphogenesis (black module).

**Figure 5.**
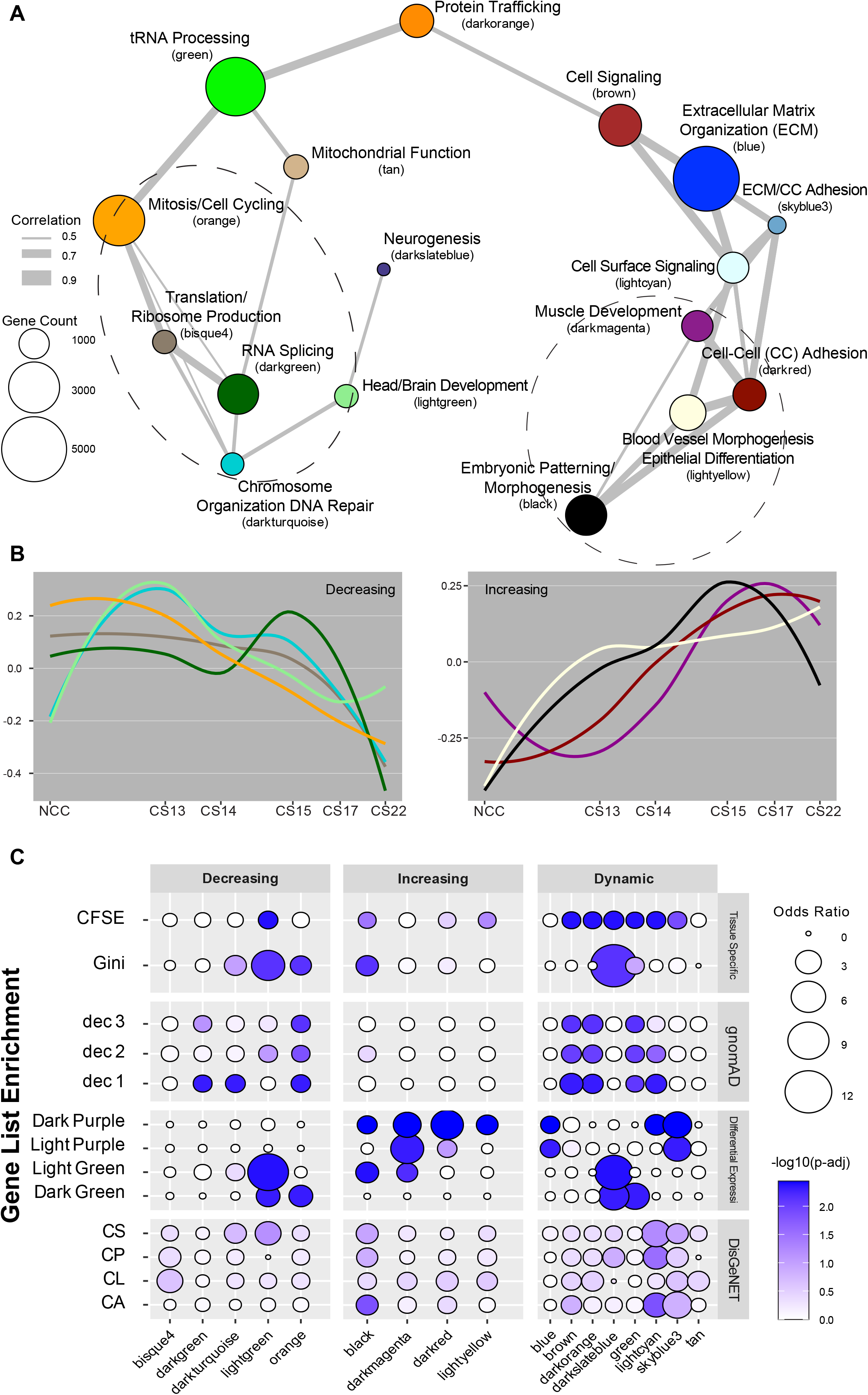
Developmental transcriptome organize into modules of co-expressed genes characterized by curated gene sets. (A) Network plot of gene modules identified by WGCNA. A Pearson correlation of the module eigenvectors was calculated for the edges where positive correlations of ≥0.5 were included. The location of each module is determined by multidimensional scaling (MDS) of the module eigengene vectors. Modules are color coded based on names assigned by WGCNA. Size of dots indicate the number of genes in each module. Each module is labeled based on the most significant biological process category gene ontology enrichment determined by DAVID; however, this label is not always all encompassing. See Table S5 for exhaustive list. Modules were grouped based on proximity and eigengene trajectory over developmental time. (B) Trajectories of expression based on eigengene vectors reported by WGCNA for each module across the developmental time series. The modules are assigned to one of three groups based on if it is generally decreasing, increasing or oscillating over developmental time. See Figure S10 for all module trajectories. (C) Dot plots of gene list enrichment within the WGCNA modules. Gene lists (rows) are divided into four groups based on their source. Dec 1 through dec 3 are genes from these deciles as reported by gnomAD. Disease gene lists from DisGeNET (Piñero et al., 2020), CS: craniosynostosis, CP: Cleft palate, CL: Cleft lip, CA: Craniofacial abnormalities. The rest of the lists are generated from data in this study. The dot color is determined by the Benjamini & Hochberg adjusted permutation p-value (see Methods) with darker indicating higher significance. Modules (columns) are grouped by their trajectory assignment from B.

We then calculated representative gene expression for each module, or module eigengene expression, and inter-module eigengenes correlation and relatedness. Modules located near one another in multidimensional scaling space (MDS) (Figure 5A) revealed three trajectories of expression across development: increasing, decreasing, or dynamic (Figure 5B, S10).

To get a finer characterization of these modules, we determined enrichment from twelve gene lists including craniofacial related abnormalities, specificity of expression, assignment of CFSE to target genes, clusters of differential expression, and measures of resistance to deleterious mutations (Figure 5C). We found that 2 modules had enrichment of known CF disease genes and 4 modules were enriched for CF Gini genes. When we analyzed modules for potential regulation by CFSEs we found 8 modules were enriched and were heavily biased toward modules with dynamic patterns of gene expression. Three modules, lightgreen, black and darkslateblue had enrichment for both CF Gini and CFSE, further establishing a connection between enhancer activity and gene expression. The coexpression networks also captured relevant gene dynamics since many of the modules showed enrichment for the DEGs from Figure 4.

We next turned to whole exome and whole genome sequences of healthy adults collected by gnomAD, which has developed a score based on observed versus expected numbers of loss of function mutations across these data (loss-of-function observed/expected upper bound fraction-LOEUF). Genes identified in the upper deciles of this scoring metric have been previously predicted to most likely cause disease in humans (Karczewski et al., 2020). We found that 8 WGCNA modules had significant enrichment for the upper LOEUF deciles. Many of these same modules were also enriched for both known disease genes and CF Gini genes.

Coexpression of genes implies that they may be more likely to interact at the protein level. For instance, genes that encode for proteins that are members of the same complex or biological process would need to be expressed within similar points of development. So we asked how our coexpression networks might reflect protein-protein interactions (PPI). We found that indeed our WGCNA network was strongly enriched throughout for known and predicted protein protein interactions. Overall, these findings strongly suggested that co-expression networks reflect the existing biology in craniofacial development (Figure 6A).

**Figure 6.**
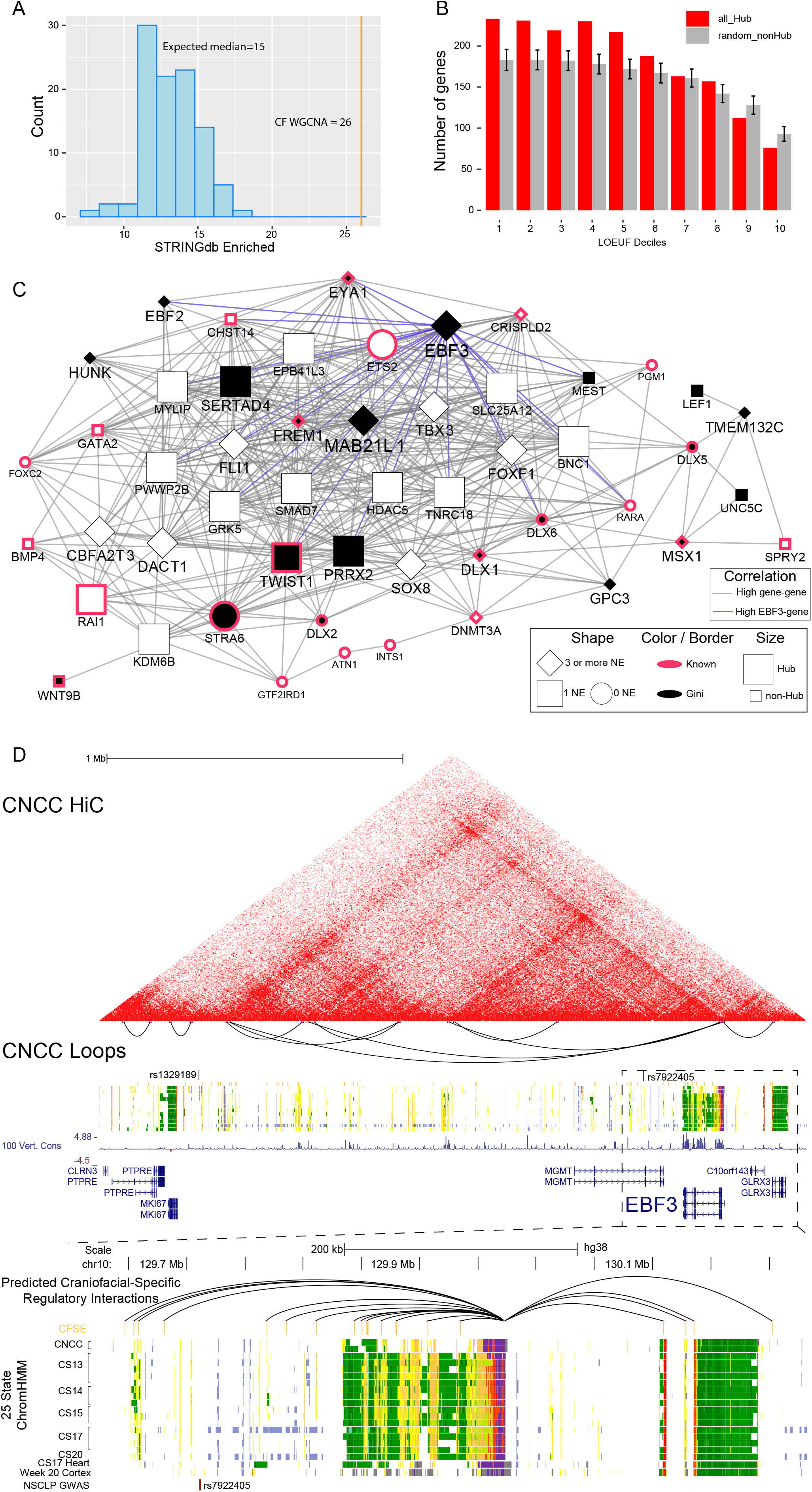
WGCNA network hub-gene enrichment of variant-restricted genes uncover disease risk of EBF3 for craniofacial development. (A) Histogram of the number of gene-scrambled modules that have protein-protein interaction (ppi) enrichment at a Bonferroni-adjusted p-value <0.05. The vertical orange line marks the number of modules that have significant ppi in the observed WGCNA network. (B) Histogram of (LOEUF) deciles of hub genes and randomly selected non-hub genes from all modules in the WGCNA network. Deciles range from decile 1 (d1), which represent the most constrained genes to d10, genes that are the most tolerant to putative loss-of-function (pLoF) variation. Error bars indicate standard deviation of 1000 permutations.(C) Network showing some of the top scoring genes (Table S6) in the black module along with known disease genes. The edges represent pearson correlation (>0.9) between genes. Blue edges connect genes that are highly correlated to EBF3. Genes are represented by nodes which are characterized by shape, fill color, border color, and size. A black fill indicates the gene has craniofacial-specific expression (Gini >=0.5). Known disease genes are marked with a fuschia border. If the gene has been assigned 3 or more CFSEs it has a diamond shape, 1 CFSE is square and 0 are circles. A node with a large size is a hub gene. Genes which have several of these criterias are displayed with larger and bold text. (D) CNCC Hi-C interaction plot of the EBF3 locus. Below is a zoomed in genome browser shot of *EBF3* locus showing 25 state ChromHMM segmentation from culture model of CNCCs, primary embryonic CF tissue, embryonic heart and fetal cortex.and CS13 through CS20 of primary human craniofacial tissue.

### Regulatory hubs in co-expression networks reveal putative driver genes for the developing face

Given the robust enrichment of multiple aspects of CF biology across multiple data types in the modules of our WGCNA, we then set out to determine if these modules could reveal novel genes involved in CF development or cause disease that had been previously inaccessible. Within coexpression networks, potential driver genes or “hub genes” are those that have correlations with many other genes (high connectivity). Previous analysis of coexpression networks of early brain and heart development showed hub genes are enriched for genes with strong evidence of involvement in ASD and congenital heart disease (CHD) risk (Cotney et al., 2015; Parikshak et al., 2013; VanOudenhove et al., 2020). We thus hypothesized that hub genes in particular would be enriched for known and novel candidate disease genes.

We first tested all hub genes against the gnomAD database and observed significant enrichment across upper LOEUF deciles and significant depletion of lower LOEUF deciles (Figure 6B). When we examined the hub genes of each module independently we identified 7 modules that were enriched across the two upper LOEUF deciles. Of these, the darkturquoise, darkred, and black modules were enriched across the top four deciles and had the highest fold enrichments over background permutations. When we combined these enrichments with the other annotation analyses and gene ontology enrichments performed above we found the black module was singularly enriched for all these metrics.

Specifically the black module was significantly enriched for genes linked to craniofacial diseases, involved in embryonic morphogenesis, CF Gini genes, genes targeted by CFSEs, constrained in healthy humans, and differentially expressed across the trajectory. We identified a number of driver genes within the black module that are well known to the CF development field including *DLX5, DLX6, EYA1, MSX1, PRRX2, MSX1* and *TWIST1*. We inspected all modules from the WGCNA that had the following traits: CF Gini genes, CFSE target genes, hub genes and high LOEUF (decile <= 3) genes. There were 539 genes that fulfilled multiple criteria, 268 of which were not in the DisGeNET database (Tables S6,S7). We consider these genes novel or previously unappreciated craniofacial disease genes warranting further study.

To demonstrate how our data can be used to prioritize and further investigate candidate disease genes, we examined *MAB21L1* and *EBF3* from the black module (Figure 6C,D). These genes shared features of known CF disease genes in the black modules, but were more highly connected, targeted by highest numbers of CFSEs, specifically expressed in CF tissues, and not widely known to play roles in CF development or disease based on annotations in OMIM and DisGeNET. *MAB21L1* has recently been identified to be responsible for an ultra rare disorder which notably involves ocular anomalies, facial dysmorphism and microcephaly (Rad et al., 2019). *Mab21l1* KO mice have ocular anomalies and lack of closure of the anterior fontanelle (Nguyen et al., 2017).Homozygous *Ebf3* mutant mice die perinatally but do not have overt CF defects. (Jin et al., 2014) Heterozygous *Ebf3* mutant mice have defects in olfactory neuron projection consistent with reported patterns of *Ebf3* expression being strongest in the olfactory epithelium (Wang et al., 1997, 2004). Only a small number of human patients have been identified with rare de novo mutations in *EBF3* but these patients consistently displayed intellectual disability, ataxia, and facial dysmorphologies (Blackburn et al., 2017; Chao et al., 2017; Harms et al., 2017; Lopes et al., 2017; Sleven et al., 2017; Tanaka et al., 2017).

Again our findings would support a role of *EBF3* in CF development, but the distinct differences in mouse and human phenotypes suggest this gene may have different functions or expression patterns across species. We also find that an ortholog of *EBF3, EBF2* is highly coexpressed (Figure 6C) consistent with previous studies showing these genes overlap in expression and function (Chiara et al., 2012; Garel et al., 1997) suggesting that this gene family may play an unappreciated role in human CF development that may not be conserved in the mouse.

### Single-nuclei RNA-sequencing of embryonic face

Given the prominence of *EBF3* in our multi-layered analyses and differences in phenotypes in human and mice harboring mutant and loss of function mutations, we hypothesized that this gene and its ortholog *EBF2* might be expressed in different locations or cell types in human craniofacial development compared with mouse. To address this question we performed single-nuclei RNA-seq on two CS20 (~11 pcw) samples. We identified 6 major cell populations (Mesenchyme, Ectoderm, Endoderm, Muscle progenitor cells, Blood cells, and Neural cells) in the human data consistent with cell types originally described in the developing mouse CF tissue (Li et al., 2019) (Figure 7A, Table S8). The relative numbers of each cell type were remarkably similar across species despite potential heterochrony (Figure S11A). While these results were encouraging they were largely qualitative and could be driven by completely different sets of genes. To verify our cell types we interrogated human clusters for expression of the mouse cell type marker genes. We observed a strong concordance between analogous cell types across species (Figure S11B).

**Figure 7.**
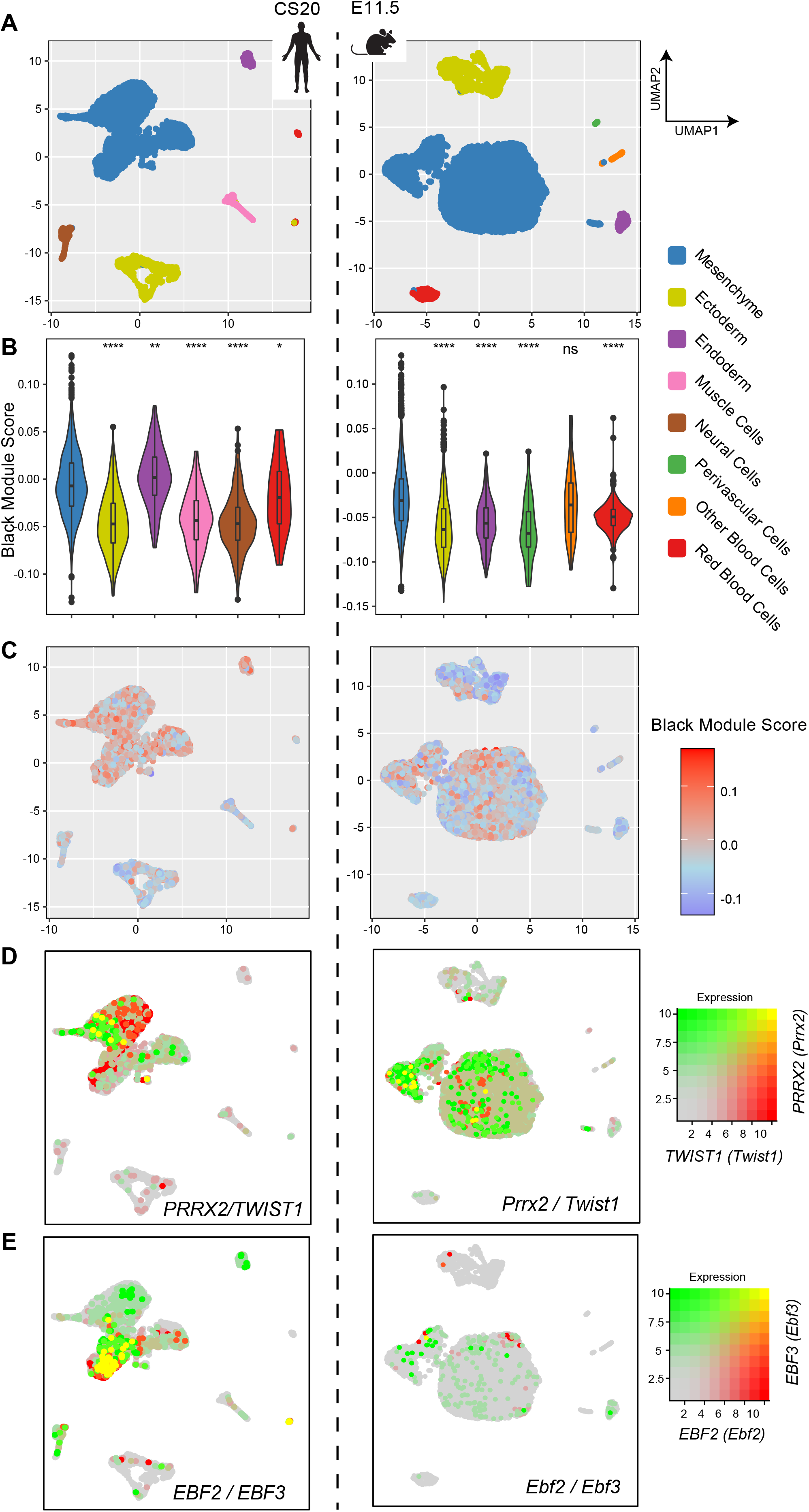
Single-nuclei RNA-sequencing of developing human face show black module enrich for mesenchyme cell populations. (A) UMAPs of CS20 and E11.5 craniofacial tissues show 6 cell populations for each. (C,D) Feature plots of top-scoring genes within the black module show enrichment of expression among the mesenchyme clusters in the UMAPs for both human and mouse and quantified in their corresponding violin plots below. Significance determined using the Mann-Whitney test (p-value ≤ 0.05 = *, ≤ 0.01= **, ≤ 0.001= ***, ≤ 0.0001 = ****) (E,F) Coexpression plots for *TWIST1* & *PRRX2* show expression dominating in the mesenchyme for both mouse and human. Green indicates high *PRRX2* (*Prrx2*) expression, red is high *TWIST1* (*Twist1*) expression and yellow is high for both, gray is low for both. (G,H) *EBF3* & *EBF2* show overlapping expression for human but not mouse mesenchyme.

As discussed above WGCNA is able to identify co-expressed genes primarily expressed in specific cell types (Parikshak et al., 2013; Rexach et al., 2020; VanOudenhove et al., 2020). To identify these gene signatures in our modules, we considered hub genes from each WGCNA module as a collective group and calculated a “module score” based upon the combinatorial activity of these groups of genes in each cell. These gene module enrichment scores were indeed enriched for specific cell types in the human data (Figure S9C). For example the hub genes of the black module showed enrichment for the mesenchyme and endoderm human cell populations. We then calculated the module score using 1:1 orthologs of genes from the black module on the mouse data (Figures 7B-C). We observed similar enrichment primarily in the mesenchyme population for mouse. When we examined expression of two known CF hub genes from the black module, *TWIST1* and *PRRX2*, we found similar patterns of expression in mesenchyme in both species (Figure 7D). We observed similar trends in cell-type restricted expression for *MAB21L1* across species again, consistent with similar roles for this gene across species (Figure S9D). However, when we considered *EBF2* and *EBF3* we observed striking differences across species (Figure 7E). We found strong coexpression at the individual cell level amongst mesenchymal cells in human consistent with the coexpression pattern in our WGCNA analysis. In contrast, we detected very little expression of either gene in mesenchymal cells of the developing mouse face and virtually no coexpression at the single cell level. When we inspected *in situ* hybridization experiments performed on E13.5 mouse embryos by the Allen Developing Mouse Brain Atlas (Lein et al., 2007) we qualitatively confirmed the patterns observed in the scRNA-Seq data. Specifically *TWIST1* and *PRRX2* were expressed in many of the same locations throughout the CF region, particularly in mesenchyme derived structures. *EBF3* on the other hand was largely absent from palatal mesenchyme and concentrated in the olfactory epithelium and the developing hindbrain consistent with phenotypes observed in knockout mice (Wang et al., 1997, 2004) (Figure S11E). These results suggested these two genes have been either removed from the CF mesenchymal lineage expression program in mice or have potentially been gained in humans. These findings warrant further exploration and demonstrate how our data can identify potentially important developmental genes that cannot be modeled in mice.

## Discussion

Craniofacial malformations caused by genetic perturbations in craniofacial development are one of the largest classes of human birth defects however the gene expression patterns active in these tissues and structures have until now been largely uncharacterized. Here we have described the most comprehensive analysis to date of gene expression from rare human embryonic craniofacial samples. We have performed large scale comparisons of gene expression from other tissues across the body and developmental stages. In order to prioritize and identify novel driver genes and regulatory networks we have subsequently integrated these data with other types of functional genomics data including ChIP-seq, Hi-C, and single cell RNA-Seq from embryonic stages of CF development.

We demonstrated that the developing CF transcriptome is it’s own distinct tissue by performing comparisons of our CF samples to thousands of samples across dozens of tissues collected by the GTEx consortium as well as developing heart and brain. Employment of an unbiased measure of specificity, the Gini index, has identified many genes that are expressed exclusively and most strongly in these novel samples. These CF Gini genes were enriched for genes known to be involved in CF disorders in humans and mice (Figure 2B), and included many genes that were either not known to be expressed in CF development or not appreciated to have highly restricted expression. By integration with our previously described (Wilderman et al., 2018) chromatin state segmentations we demonstrated that these specifically expressed genes are systematically surrounded by craniofacial-specific enhancers over distances of ~400kb. These genes represent excellent disease candidates and warrant further exploration by the field (Table S7).

Analysis of time-series differential gene expression revealed dynamic gene utilization that coincided with known morphological changes throughout development. Our work identified two subsets of dynamic genes that are enriched for craniofacial disease ontologies. Their trajectories of expression peak during 4 to 4.5 post-conception weeks (CS13-CS14) and again at 5.5 to 6 post-conception weeks (CS15-CS17) (Figure 4C). The fusing of major facial prominences is believed to occur during this time period, the failure of which leads to CL/CLP. These pairwise analyses are crucial for helping to understand how gene expression changes time, however many genes that are important for craniofacial development with steady expression across our samples may be overlooked. A prominent example is *MSX2* which has robust expression in our data and is identified by the Gini index, but does not rise to significance across the time series. To address this issue we employed coexpression analysis with a commonly used approach, WGCNA. WGCNA has been demonstrated to identify biologically relevant gene networks in a variety of contexts and allow for more nuanced complex gene expression dynamics to be detected.

The WGCNA network we constructed has several modules significantly enriched with genes involved in embryonic patterning, cell-signaling, and cell-cell adhesion (Figure 5A). More specifically we found two modules have enrichment for known craniofacial disease genes and one of these, the black module, is enriched for CF Gini genes and CFSE targeted genes. These findings prompted us to examine this module more closely for novel candidate disease genes. In particular genes that are coexpressed with many other genes or potential “hubs” of regulation have been shown to be critical for model organisms (Albert and Barabási, 2002; Barabási and Oltvai, 2004; Carlson et al., 2006).

Highly connected hub genes from coexpression analysis are also likely to be mutation resistant in humans, and thereby excellent candidates for disease. We showed that genes with constrained mutation in the human population are enriched in the hub genes within our coexpression networks (Figure 6A). However it’s important to note that not all identified hub genes are relevant within the system (Langfelder et al., 2013) so it is necessary to integrate other layers of data to identify candidates genes of craniofacial disease specifically. We addressed this by incorporating craniofacial specific expression and putative craniofacial specific enhancer targeting to look across all modules for these characteristics shared with the genes in the black module. In particular we observed that the lightgreen and darkslateblue modules contain many craniofacial relevant hub genes such as those in the *WNT* family and *PHOX2* genes. This allowed us to unbiasedly identify 540 genes, 239 of which are not known to be associated with human craniofacial disease. This list of genes and the craniofacial regulatory regions that surround them are excellent candidates for further study. From the black module we examined the EBF3 locus chromatin architecture and cell-specific expression to demonstrate how to leverage this resource.

EBF3 has been associated with neurodevelopmental problems along with craniofacial dysmorphism. There are 2 GWAS variants for orofacial clefting located in enhancers at the locus (Chen et al., 2018; Leslie et al., 2016; Shi et al., 2012). One is located in an intronic region of MGMT and therefore was predicted to affect that gene, however MGMT is lowly expressed in our data, and we contend that EBF3 is a better candidate due to its conspicuous signals from multiple genomic data types in our analysis.

Co-expression networks have been shown to deconvolute cell-type gene signatures from bulk RNA-seq data, implying that cell types exclusively utilize sets of coexpressed genes for establishing cell identity. Thus we reasoned that genes from the black module might be enriched in specific cell types in the developing mammalian face. Indeed when we performed snRNA-Seq on primary human tissue and jointly analyzed previously published mouse scRNA-Seq data we found that genes from the black module are enriched for mesenchyme cell markers in both human and mouse. When we inspected these single-cell data sets for well known craniofacial genes of the black module like TWIST1, we found similar patterns of cell restricted expression between human and mouse (Figure 7D). However when we examined EBF3 and EBF2 we observed a striking difference in absolute level of expression in the mesenchyme and co-expression in the same cells between the two species. These two genes were strongly coexpressed in a subset of mesenchyme cells in human, but virtually absent in mouse (Figure 7E). This could be due to differences in heterochrony in embryos across species. However when we examined gene expression in publicly available in situ hybridization data from the E13.5 mouse, a time point where the developing mouse face is morphologically closer to CS20, we observed EBF3 expression in the developing olfactory system and cerebellum but virtually absent from mesenchyme derived tissues (Supplemental S11E). These expression patterns are consistent with the phenotypes observed in mouse knockout models (Wang et al., 1997, 2004) and raise the possibility that either the EBF family of genes has gained new activities in craniofacial development on the primate lineage or rodents no longer employ them in this way. These findings also illustrate the need for more gene expression data from the embryonic period of human development and comparative approaches across species. Many models of disease genes are verified in mouse, however this may lead to overlooking important genes that are exclusively utilized by humans.

In closing, by integrating multiple layers of functional genomics data including gene expression, chromatin states, chromatin structure and population genetics, we have provided compelling evidence for novel disease genes in human craniofacial development. We provide all data in formats that are directly comparable to other large consortia, and can be downloaded from Gene Expression Omnibus. We have updated our existing public track hubs on the UCSC Genome Browser and the Cotney Lab website. For ease of data exploration, we have provided the bulk and single cell data in this study as interactive shiny applications http://cotneyweb.cam.uchc.edu/craniofacial_bulkrna/ and http://cotneyweb.cam.uchc.edu/craniofacial_cs20/. We hope these resources can be used by the field to better understand human craniofacial development and aid clinicians in interpreting genetic data from craniofacial abnormality patients.

## Acknowledgements

We would like to thank members of the UConn/JAXGM Single Cell Genomics Core for help with standardizing single cell isolation techniques and preparing of sequencing libraries. We would also like to thank members of the UConn Computational Biology Core and High Performance Computing Facility for assistance with package installation and software/hardware support. This work was funded by grants from the National Institutes of Health NIDCR 1R01DE028945, NIDCR 1R03DE028588, and NIGMS 5R35GM119465.

## Author Contributions

Conceptualization TNY and JC. Investigation AW, EW, and JV. Formal Analysis TNY, EW, and JC. Writing - Original Draft TNY and JC. Writing - Review and Editing, TNY, AW, JV, and JC. Funding Acquisition JC. Supervision JC.

## STAR METHODS

### RESOURCE AVAILABILITY

#### Lead contact

Please direct any further requests for resources and code to Dr. Justin Cotney.

#### Materials availability

This work did not generate any new or unique reagents.

#### Data and code availability

- Data can be interactively explored at http://cotneyweb.cam.uchc.edu/craniofacial_cs20/ and http://cotneyweb.cam.uchc.edu/craniofacial_bulkrna/.
- All original code has been deposited in the cotney lab github (https://github.com/cotneylab) and is publicly available.
- Any additional information required to reanalyze the data reported in this paper is available from the lead contact upon request.

## EXPERIMENTAL MODEL AND SUBJECT DETAILS

### METHOD DETAILS

#### Human Tissue Samples

Use of human embryonic tissue was reviewed and approved by the Human Subjects Protection Program at UConn Health (UCHC 710-2-13-14-03). Human craniofacial tissue was collected, staged, and provided by the Joint Medical Research Council (MRC)/Wellcome Trust Human Developmental Biology Resource (www.hdbr.org). Tissues were flash frozen upon collection and stored at −80°C.

#### RNA-seq

Frozen tissue samples were added to Qiazol (Qiagen) and subjected to mechanical disruption using a motorized pestle. Homogenates were then processed using the miRNeasy RNA extraction kit (Qiagen, 217004) with on-column DNAse treatment (Qiagen, 79254) according to the manufacturer’s protocol. RNA integrity was checked using Agilent Tapestation 2200 with Agilent RNA analysis screentapes (Agilent Genomics, 5067-5576). RNA with RNA Integrity Number (RIN) scores preferably > 8.0 were used in the preparation of RNA-seq libraries. RNA-seq libraries were prepared from 100-200ng total RNA using the TruSeq stranded mRNA kit (Illumina, RS-122-2101) according to the manufacturer’s instructions with the modification to use Superscript III Reverse Transcriptase enzyme (Invitrogen, 18080044) during the first strand cDNA synthesis step. Completed libraries were checked for quality and average fragment size using the Agilent Tapestation 2200 with D1000 screen tapes (Agilent Genomics, (5067-5582). Molar concentration determined using NEBNext qPCR library quantification kit (NEB,E7630). Libraries were pooled and diluted to 1.8pm and sequenced on the NextSeq500 Illumina platform using 75bp paired end sequencing according to manufacturer’s recommendations. Libraries were diluted to 4nM, pooled and denatured according to the instructions for Illumina NextSeq 550/500. Libraries were sequenced on the NextSeq 500 or 550 with settings for single-index, paired-end sequencing with 75 cycles per end.

#### snRNA-seq

Samples were mechanically disrupted into liquid suspensions, checked for viability and counted using Trypan blue staining. Nuclei were isolated and quantified following the established protocol (10x Genomics^®^). Samples were transferred to the Jackson Laboratories (Farmington, CT) Single Cell Biology Laboratory (SCBL) for processing which followed the *Chromium Next GEM Single Cell Multiome ATAC + Gene Expression* user guide from 10x Genomics^®^. Sequencing was done on an Illumina NovaSeq.

### QUANTIFICATION AND STATISTICAL ANALYSIS All plots made with ggplot(mention in key resources)

#### RNA-seq Data Processing

Quality control was performed on RNA-seq reads using FastQC (v.0.11.7) and MultiQC (v.1.1) (Ewels et al., 2016). Trimming for adapters, quality and length was performed using Trimmomatic (v.0.36) (Bolger et al., 2014). Trimmed fastqs were aligned with Rail-RNA (v.0.2.4b) (Nellore et al., 2017) using human assembly GRCh38/hg38. RSeQC (v.4.0.0) (Wang et al., 2012) was used to calculate the read distribution, gene body coverage and TIN score (Figure 1B, S1A) (Wang et al., 2016). The coverage bigWig files output by Rail-RNA were used as input for the generation of counts tables by following the instructions and pipeline from *recount2* (https://github.com/leekgroup/recountcontributions), where the comprehensive gencode v.25 annotation was used. The *level 3* genes as defined by gencode were excluded. The recount rse_gene objects for each sample were combined into one rse_gene object and transformed with scale_counts() from recount (v. 1.8.2). The PCA plots in Figure 1B and S2 were made using the prcomp() function from the built-in R stats package on a DESeq2 (v.1.26.0) (Love et al., 2014) rlog() transformation on the raw counts of the 18597 most highly expressed genes across all craniofacial samples generated in this study.

#### snRNA-seq Data Processing

Raw fastqs were aligned to Gencode37 using Kallisto-Bustools (kallisto v.0.46.2, bustools v.0.40.0) (Melsted et al., 2021) and gene counts per cell were imported into R using BUSpaRse (v.1.0.0). Seurat (v.3.2.0) (Stuart et al., 2019) was used for filtering, merging of replicates, scaling, normalization, dimensionality reduction (UMAP), and clustering. Clusters were functionally annotated using GO enrichment analysis in clusterProfiler (v.3.14.3) (Wu et al., 2021; Yu et al., 2012) from marker genes of each cluster. Cluster annotations were confirmed using modules of 1:1 orthologs of marker genes from mouse craniofacial tissue cell types (Li et al., 2019). Enrichment of the black WGCNA module hub genes per cell type (Figure 7B,C) were calculated using AddModuleScore(). The coexpression UMAPs were calculated using FeaturePlot() with blend =TRUE and order=TRUE.

#### GTEx Analysis

The rse_gene R data object, or counts table of all GTEx tissues was retrieved from the recount2 database (https://jhubiostatistics.shinyapps.io/recount/). The GTEx counts table was generated using the same Rail-RNA & recount2 pipeline that was used to generate the counts tables for our embryonic craniofacial data which is described above. The GTEx dataset contained 9,662 bulk RNA-seq samples which were combined with 24 embryonic heart samples (VanOudenhove et al., 2020), and 5 fetal cortical plate samples (Fietz et al., 2012) and our 12 embryonic craniofacial samples. The metadata for GTEx is provided in a link under the phenotype column from the recount2 database, and the tissue assignments located under the column named *smts* was used, resulting in a total of 34 unique tissues. The counts were transformed using the scale_counts() function from the recount R package.

The plots in Figure 1D were analyzed using a down-sampled version of GTEx, where twelve random samples from each tissue were chosen. The counts matrix was filtered for lowly expressed genes using rowMeans() >50. A transformation was performed using vst() function from DESeq2. The function plotPCA() from DESeq2 using ntop= 500 most variable genes was used for the PCA plot. The spearman correlation between the vst-transformed down-sampled tissues was calculated and plotted in R using the pheatmap (v. 1.0.12) package.

The tSNE plot in Figure S1C was made using GTEX, embryonic heart, fetal brain, craniofacial samples counts table filtered for lowly expressed genes. The filtered counts matrix was transformed by log10() with a pseudo count of 1 added to all values. The transpose of the log10 transformed matrix was then converted to a distance matrix using the dist() function in R. This distance matrix was used as input for the tSNE model generated by using the Rtsne() function from the R package Rtsne (v.0.15). The parameters used in Rtsne() were the following: perplexity=100, max_iter=1000, theta=0.5, dims=2.

For qualitative signal comparisons across tissues in Figure 1C, the UCSC genome browser was used to load the GTEx RNA-Seq Signal track hub. Expression signals from 30 samples of each tissue type indicated were randomly selected. BigWigs generated by the Rail-RNA pipeline above for craniofacial samples were visualized alongside these tracks at the *TWIST1* gene locus.

#### Tissue Specificity of Gene Expression (GINI)

A total of 34 tissues, 31 from GTEx, and embryonic heart, craniofacial, and fetal brain was used to calculate the tissue specificity. The Gini Index for each gene was calculated using the Gini() function from R library Ineq (v0.2-13) on the average counts across all samples per tissue. A gene was given a tissue assignment based on the tissue with the maximum average count across all tissues for that gene. To create the heatmap of most restricted genes, the distance matrix of one minus the transpose of the Pearson correlation of the average expression per tissue for all genes with a Gini score of 0.7 and above was clustered using hclust() with method = “complete”. For visualization purposes, in the heatmap of Figure 2A sex tissues including Breast, Testis, Vagina, Cervix Uteri, Fallopian Tube, Prostate, Ovary and Uterus were excluded but are included in Table S1. Testis are known to have a large amount of upregulated genes thus likely contributing to the large amount of Gini genes assigned to it (Melé et al., 2015). The average expression per tissue matrix was plotted using pheatmap with scaled rows organized by the dendrogram calculated from hclust(). Disease enrichment analysis was done using the R package disgenet2r (v0.99.2) disease_enrichment() function on genes 0.7 and above. The fold enrichment score was calculated by dividing the *Ratio* by the *BgRatio* columns and taking the log base 2.

#### Novel craniofacial enhancer effects on gene expression

Assignment of enhancers to genes was made using GREAT (v.4.0.4) on craniofacial specific enhancers (CFSE) hg19 with the whole genome as background and using the single nearest gene association rule setting to 1MB. The line graph in Figure 3A of the enhancers versus expression used the average expression per tissue type from GTEx and our heart enhancer data using geom_smooth() from ggplot2 with the method set to “gam”.

#### Distance to novel craniofacial enhancers analysis

For Figure 3B We used gencode annotation v25 to get location of TSS for all genes. A Gini gene was defined as having >= 0.7 gini score. Bedtools (v.2.28.0) closest was used to get the distance between the closest gini TSS to CFSE. A background distribution was established by picking distance to random set of 239 gini genes among all tissues (n=33) excluding those assigned to CF at 1000 permutations. The interpolation lines were plotted using geom_smooth() with formula=y~log(x). To get the median closest gene-enhancer for our data, we only considered enhancer pairs within 400kb distance as that was significant over random.

Figure S3A and S3B use the Gasperini et al. 2019 data. We plotted a histogram for their 664 high-confidence enhancer gene pairs to compare directly to Figure 3B. Figure S3B is a violin plot of these same 664 high-confidence enhancer gene pairs and our CF gini-CFSE pairs that were <= 1 Mb (n=3437).

#### Differential Expression

The scaled rse_gene recount object including NCC, CS13,CS14,CS15,CS17 and CS22 samples were made into a DESeq2 (v.1.22.2) object. Low gene counts were filtered by removing all genes whose mean of counts across all samples was less than 50. This left a total of 18597 genes for downstream analysis. Batch effects were mitigated by using the sva (v.3.30.1) package in R. Included the CS13, CS14, CS15, CS17 and CS22 for the sva calculation which resulted in 4 surrogate variables. Pairwise differential analysis between CS13, CS14, CS15, CS17 were performed with DESeq2 including the surrogate variables. Differentially expressed genes were plotted in the heatmap of Figure 4A considered if they had a Benjamini Hochberg adjusted p-value of less than 0.05 and a log2 fold change of greater than absolute value of 1. The PCA plot in Figure S6A was made as described above except on just the 3677 differentially expressed genes. The R package Vennerable (v.3.1.0.9) was used to make Figure S6B.

The heatmap in Figure 4A was made using the R package pheatmap (v.1.0.12) on the rlog() transformation of the raw counts. The rlog() counts distance matrix of one minus the transpose of the kendall correlation was clustered using hclust() with method = “complete”. The counts were scaled by setting the pheatmap scale option to “row”. The resulting hierarchical clustering was used in the pheatmap option cluster_rows to organize the gene rows and the hierarchical clustering of the sample columns was from pheatmap default options. ClusterProfiler (v3.12.0) was used to obtain the gene and disease ontology enrichments from the dendrogram clusters (Figure 4B, S4E) using the function enrichGO() with standard options. The fold enrichment score was calculated by dividing the *Ratio* by the *BgRatio* columns and taking the log base 2

In Figure 4C, the disease ontologies were obtained for the lightgreen (cluster 2) differentially expressed genes using enrichDGN(). The network plot was made using cnetplot() from enrichplot (v.1.6.1) using log_2_ fold change between CS13 and CS17 from DESeq2.

The heatmap in Figure S6D was made as described above, utilizing 6 samples from GEO accession GSE70751, SRX numbers SRR2096446 through SRR2096451 linked to (Prescott et al., 2015). The fastqs were processed using the same Rail-RNA/recount pipeline as all other data in this paper.

#### Gene class analysis of differentially expressed genes

Three lists were used to represent most transcription factors (Lambert et al., 2018) (n=1639), RNA-binding proteins (Cook et al., 2011) (n=415) and genes annotating the GO biological process cell-cell signaling, GO:0007267 (n=1364). Trajectory plots in Figure S7A,B were made using geom_smooth() from ggplot2. The quantitative locations of x-axis depicting time points were determined by taking the median x-value for each timepoint from the PCA plot of Figure 1B. Enrichment over background (Figure S7C) was determined by randomly selecting 3677 genes from the 18597 used in the DESeq2 analysis 1000 times.

#### ChIP-seq data integration

##### Across differentially expressed genes

For Figure S8: Imputed chromatin signals for each histone mark and DNAse hypersensitivity were extracted in a 5kb window surrounding the transcription start sites of genes from each indicated cluster using computeMatrix reference-point command in deepTools (v.3.5.1). Histograms and heatmaps were generated using the output of computeMatrix in plotHeatmap command of deepTools.

##### Differential H3k27ac signal

For Figure S9A: H3K27ac counts at each craniofacial enhancer region were generated from bigWig files of imputed H3K27ac -log10(p-value) signals using rtracklayer and GenomicRanges in R (3.5.2) based on an original read length of 75 bp. Heatmap constructed from H3K27ac counts filtered for those determined to be significantly different between CS13 and CS17 using DESeq2 for the following cutoffs: Benjamini-Hochberg p-adjusted < 0.01 and a log_2_ Fold Change > abs(1). The counts distance matrix of one minus the transpose of the Pearson correlation was clustered using hclust() with method = “complete”. The dendrogram was cut into two clusters and the H3K27ac segments from each cluster were intersected with genome-wide TSS using bedtools (v.2.28.0) The permutation test to establish a background of intersecting genes was done by randomly selecting segments from the genome using bedtools shuffle with the -chrom setting. The genes of these TSS’s were intersected with the differentially expressed genes from Figure 4.

#### Weighted Gene CoExpression Network Analysis

We generated co-expression networks using the WGCNA R package based on recommendations put forth by the Horvath group.

##### Network Construction

The rse_gene object from the recount pipeline was combined for all samples resulting in a starting matrix of all genes (gencode v.25) by 17 samples including 3 replicates of CNCC, CS13, CS14, CS15, CS17 and 2 replicates of CS22, the raw counts were scaled by using scale_counts() from recount (v1.8.2). Lowly expressed genes were filtered by excluding all genes whose sum across all samples were lower than 100. SVA was used to transform the counts to account for batch effects, 4 surrogate variables were detected. All resulting negative counts were set to 0. The counts matrix was further transformed by log_2_ with a pseudo count of 1.

A soft-thresholding power of 18 was chosen assuming a signed network and based on recommendations for less than 20 samples. The modules were detected from the network from the cutreeDynamic() function from the WGCNA package with the following parameters, minClusterSize=100, deepSplit=2. Detected modules were merged based on their eigengene correlation. To do this a dendrogram of the module eigengenes was generated and a threshold value of 0.18 was chosen as input for the function mergeCloseModules(). The resulting network contained 29 modules.The intra-modular connectivity of each gene was calculated using the intramodularConnectivity() function from the WGCNA library which was the metric used to determine the hub and non-hub designation.

##### Plotting of Modules

A multidimensional scaling of the module eigenvectors output from WGCNA was generated to plot the modules in 2-D space using the function cmdscale() from the stats (v.3.6.1) R package. A Pearson correlation of the module eigenvectors was calculated for the edges. Positive correlations of 0.5 and greater were included. Modules were plotted that fulfilled the criteria of having significant adjusted p-values (< 0.05) from the GO analysis, or significant (Benjamini & Hochberg) adjusted permutation p-values (< 0.05) in at least two of the gene lists from the Tissue Specific, gnomAD, or DisGeNET categories (Figure 5C). The module eigenvectors of each of the 17 modules were plotted using the geom_smooth() function with method= “loess”. The quantitative locations of x-axis depicting time points were determined by taking the median x-value for each timepoint from the PCA plot of Figure 1B. The confidence intervals were removed for ease of visualization. Figure 6C was made using the open source software Cytoscape (v.3.8.2) (Shannon et al., 2003).

##### Gene Ontology and Functional Enrichments

RDAVIDWebService (v1.22.0) was used to obtain gene ontology enrichment of the genes within each of the 29 WGCNA modules. The gene background list used was all the genes input into the WGCNA. The module enrichments of gene lists were determined by a permutation test. The 26626 genes were randomly scrambled among the same number and size of modules as the original network and overlapped with the respective gene lists. This was repeated for 1000 iterations to calculate a permutation p-value adjusted by (Benjamini & Hochberg).

##### Protein-protein interaction analysis

To generate the ppi histogram (Figure 6A), 100 randomized versions of the WGCNA network were made. This was done by randomly assigning the 26626 genes to 29 modules of equal gene sizes to the original network using the R function sample(). The ppi enrichment of up to 500 randomly chosen genes for each module of each of the 100 randomized versions was then determined using the STRINGdb (v1.24.0) package. Up to 500 genes were used due to constraints from the STRINGdb package. The ppi database was loaded by using STRINGdb$new with version =“10” and score_threshold=0.4. For each iteration, the output p-value of the STRINGdb call get_ppi_enrichment() was adjusted using the Bonferroni method. The number of modules that met the adjusted p-value cut off of 0.05 was counted for each iteration to produce frequency values.

##### Hub gene LOEUF enrichment

For the LOEUF enrichment analysis among the hub vs non-hub genes (Figure 6B), the non-hub genes were randomly sampled using the R function sample(). The number of non-hub genes sampled were the same number of total hub genes within the network, the 10% with highest connectivity of 26626 genes or 2,663 from the non-hub gene list. This process was iterated 1000 times to get a mean and standard deviation for each LOEUF decile. The LOEUF score and decile designation for each gene is freely available through gnomAD v2.1.1

##### Novel gene list

The following parameters were used to generate Table S7. The gene must be a CF Gini gene AND one of the two: CFSE targeted OR a hub gene, AND have a LOEUF decile score of <= 3 OR be a known CF disease gene within the four categories from DisGeNET (Piñero et al., 2020), craniosynostosis, Cleft palate, Cleft lip, Craniofacial abnormalities.

## Supplemental Information Titles and Legends

**Table S1.** Embryo metadata. Metadata as reported by HDBR for embryonic sequencing samples generated by our lab (Columns 1 through 6). RIN scores determined in this study, batch numbers represent separate individuals generating the libraries.

| Replicate ID | Sample ID | Carnegie Stage | Karyotype | Collection Site | Collection Date | Batch | RIN score | Experiment |
| --- | --- | --- | --- | --- | --- | --- | --- | --- |
| R2 | 13365 | CS13 | 46_XY | London | 09/27/16 | 2 | 8.4 | Bulk RNA-seq |
| R3 | 13472 | CS13 | 46_XY | London | 12/06/16 | 2 | 8.6 | Bulk RNA-seq |
| R1 | 12690 | CS13 | 46_XY | London | 06/23/15 | 1 | 9.1 | Bulk RNA-seq |
| R3 | 12408 | CS14 | 46_XY | London | 01/20/15 | 1 | 8.4 | Bulk RNA-seq |
| R1 | 12704 | CS14 | 46_XY | London | 06/30/15 | 1 | 7.6 | Bulk RNA-seq |
| R2 | 12913 | CS14 | 46_XX | London | 11/03/15 | 1 | 8.9 | Bulk RNA-seq |
| R3 | 13128 | CS15 | 46_XY | Newcastle | 06/29/16 | 1 | 9.1 | Bulk RNA-seq |
| R2 | 13000 | CS15 | 46_XX | Newcastle | 06/29/15 | 1 | 6.7 | Bulk RNA-seq |
| R1 | 13019 | CS15 | Unknown | Newcastle | 06/29/16 | 1 | 8.9 | Bulk RNA-seq |
| R2 | 12320 | CS17 | 46_XY | Newcastle | 12/07/15 | 1 | 7.4 | Bulk RNA-seq |
| R3 | 12421 | CS17 | 46_XY | Newcastle | 12/07/15 | 1 | 9 | Bulk RNA-seq |
| R1 | 12588 | CS17 | 46_XX | Newcastle | 12/07/15 | 1 | 8.4 | Bulk RNA-seq |
| R1 | 13327 | CS20 | 46_XX | Newcastle | 08/26/16 | N/A | N/A | Single Cell Multiome |
| R2 | 13503 | CS20 | 46_XY | Newcastle | 01/19/17 | N/A | N/A | Single Cell Multiome |

**Table S2. List of Gini scores and tissue specificity for each gene.** Gini score (0 to 1) for 33073 genes across 34 tissues.

**Table S3. Bed file of craniofacial specific enhancers.** Regions determined by chromatin state segmentation (ChromHMM) and include states 13,14,15 and 18. The second tab contains results from GREAT for these segments (McLean et al., 2010).

**Table S4. Pairwise differential expression results.** Results from DESeq2 with raw scaled counts tables attached. The last tab contains information for each gene and their cluster designation. Related to Figure 4.

**Table S5. Gene ontology results of each WGCNA module.** Related to Figure 5A.

**Table S6. WGCNA results.** A list of genes assigned to each module separated by tabs. Integrative information included such as Gini status and known disease link. Related to Figure 5 and 6.

**Table S7. Novel CF disease candidate list.** *Designation* column indicates if gene is in the DisGeNET database for craniofacial-related diseases: Craniosynostosis, Cleft palate, Cleft lip, Craniofacial abnormalities.

**Table S8. Single-Nuclei RNA-seq marker genes.** Results output from Seurat. Related to Figure 7.

**Figure S1. Developing craniofacial tissue quality control metrics and global multi-tissue comparison.**

**A.** Read distribution plot generated by RSeQC. PolyA RNA samples dominated by CDS_exons while the two total-RNA samples (CS22) dominated by Intronic regions as expected. **B.** PCA plots of global gene expression for bulk RNA primary tissue samples generated in this study. The left indicates CS staging. The right plot marks karyotyping and RIN values for each sample, XX (female), XY (male), UNK (unknown). **C.** Comparison of raw counts from GTEx with early developing tissues. The top plot is the number of genes above 10 counts per sample per tissue. The bottom plot is the gene expression ranges for the average expression across all samples per tissue. **D.** tSNE plot of global gene expression counts for multi-tissue analysis (craniofacial, embryonic heart, fetal brain, GTEx). Representative tissues are highlighted by color: craniofacial (blue), embryonic heart (red), fetal brain (light green), GTEx brain (green), GTEx Heart (purple), GTEx muscle (yellow), all other GTEx tissues (grey).

**E.** Violin plots of TWIST1 expression across all samples per tissue surveyed. Related to Figure 1C. Stat_compare_means() was used to compare all tissues against the craniofacial tissue, using the Mann-Whitney test (p-value ≤ 0.05 = *, ≤ 0.01= **, ≤ 0.001= ***, ≤ 0.0001 = ****)

**Figure S2. Signaling-related gene ontologies of craniofacial Gini genes.**

Selected set of significant (Benjamini and Hochberg–adjusted p-value <0.05) gene ontologies for gini genes that involved signaling. The dot size or Count indicates number of genes within a disease category. Color bars are log2 of the fold enrichment (see Methods).

**Figure S3. Gene-enhancer pair distances are larger for developing tissue. A.** Histogram plot of distances for high-confidence gene-enhancer pairs for K562 cells from Gasperini et al. 2019. **B.** Violin plots comparing the distances for our CF Gini gene-CFSE pairs to Gasperini data (see Methods). **C.** Gini gene enrichment across all tissues for all genes with bivalent promoters in craniofacial tissue. The median and standard deviation were calculated from 1000 iterations of randomly selected genes.

**Figure S4. Developmental craniofacial transcription factors genome browser shots.** Genome browser shot of gene locus showing 25 state ChromHMM segmentation from culture model of CNCCs, and CS13 through CS20 of primary human craniofacial tissue. The dark purple indicates bivalent state which are located near or at the promoter (reds). Above the segmentations track is the active and novel CFSE track shown in orange which include strong enhancer ChromHMM states 13-15. Below are RNA-seq bigwig signal tracks for human embryonic stem cells (ESC), CNCCs, and CS13 through CS22. *Top* NR2F1 *middle* MSX2 *bottom* ALX4.

**Figure S5. Novel developmental craniofacial transcription factors genome browser shots.** Genome browser shot of gene locus showing 25 state ChromHMM segmentation from culture model of CNCCs, and CS13 through CS20 of primary human craniofacial tissue. The dark purple indicates bivalent state which are located near or at the promoter (reds). Above the segmentations track is the active and novel CFSE track shown in orange which include strong enhancer ChromHMM states 13-15. Below are RNA-seq bigwig signal tracks for human embryonic stem cells (ESC), CNCCs, and CS13 through CS22. *Top* BCL11A *bottom* HMGA2.

**Figure S6. Differential expression of early developing craniofacial tissues.**

**A.** PCA plot of RNA-seq samples generated in this study including only differentially expressed genes. **B.** Venn diagram of differentially expressed genes**. C.** Bar plots showing number of up and down regulated differentially expressed genes compared to each time point. For example CS17 has 2000 up regulated and 1000 down regulated genes in pairwise comparisons with CS13, CS14 and CS15. **D.** Heatmap of differentially expressed genes (rows) separated into three panels. The first panel showcases the data from (Prescott et al., 2015). The other panels are separating samples into early (CNCC, CS13,CS14) and late (CS15, CS17, CS22). Related to Figure 4A. **E.** Gene ontology of a select subset of the most significant terms identified by clusterProfiler for 1-Dark Green and 3-Light Purple. The genes belonging to 1-Dark Green are enriched for functions relating to neurogenesis and somitogenesis. The light purple has a focus on cell-cell adhesion and sarcomere organization. Related to Figure 4B.

**Figure S7. Gene class analysis of differentially expressed genes. A.** Trajectories of the z-score of the raw counts of all differentially expressed genes separated by class, TF or C2C (cell-cell signaling) and dendrogram cluster from Figure 4A. **B.** Trajectories of the z-score of the raw counts of single genes colored by the dendrogram cluster from Figure 4A. Genes in Dark Purple-4 have two separate plots to highlight differences. **C.** Bar plots of expected versus observed number of genes from each gene class. Median and standard deviation for expected values calculated from 1000 permutations. **D.** Histogram of Gini scores of all RBPs (n=415).

**Figure S8. ChIP-seq signal across differentially expressed genes from multiple histone modifications.** The ChIP data is from CS13 human primary tissue. Note that genes of the Dark Green-1 cluster have the highest signal from H3K4me3, an epigenetic marker for activation of transcription. These genes were also shown to have the highest expression in our early tissues (Figure 4A).

**Figure S9. Differential H3K27ac signal between CS13 and CS17. A.** Heatmap of H3K27ac segments (n=19010) separated into two clusters through hierarchical clustering. The Z-score indicates signal intensity where yellow is high and blue is low. Cluster 1 tends to have higher signal in the early developmental stages and Cluster 2 has higher signal in the later developmental stage (CS17). **B.** The intersection of differentially expressed genes from the clusters in Figure 4A and genes obtained by assigning H3K27ac segments using GREAT. Cluster 1 is more enriched for genes upregulated in the early stages (Dark Green-1) and cluster 2 for late stages (Dark Purple-4).

**Figure S10. Trajectory plots for WGCNA module eigengenes.** Related to Figure 5B.

**Figure S11. Single-nuclei RNA-seq on human and mouse craniofacial tissue. A.** Relative proportions of cell types. **B.** Dot plot visualization as calculated by Seurat. Comparison of marker genes (species orthologous) and their expression between human and mouse cells. **% *Expressed*** indicates percent of cells within a designated population expressing the marker genes. Color bar represents scaled average expressed level across all cells within a population. **C.** Dot plot visualization as calculated by Seurat. Expression of WGCNA module hub genes (columns) within a cell population (rows). **D.** Feature plot of *MAB21L1* and *Mab21l1* expression in human and mouse cells. **E.** In situ-hybridization (ISH) RNA probe assays in E13.5 mouse. Allen Developing Mouse Brain Atlas, *Twist1:* http://developingmouse.brain-map.org/experiment/show/100076491 *Prrx2:* http://developingmouse.brain-map.org/experiment/show/100094084 *Ebf3:* https://developingmouse.brain-map.org/experiment/show/100093321.

