## Supplementary_Figures for "Integrative analysis of transcriptomics in human craniofacial development reveals novel candidate disease genes"

### Figure S1

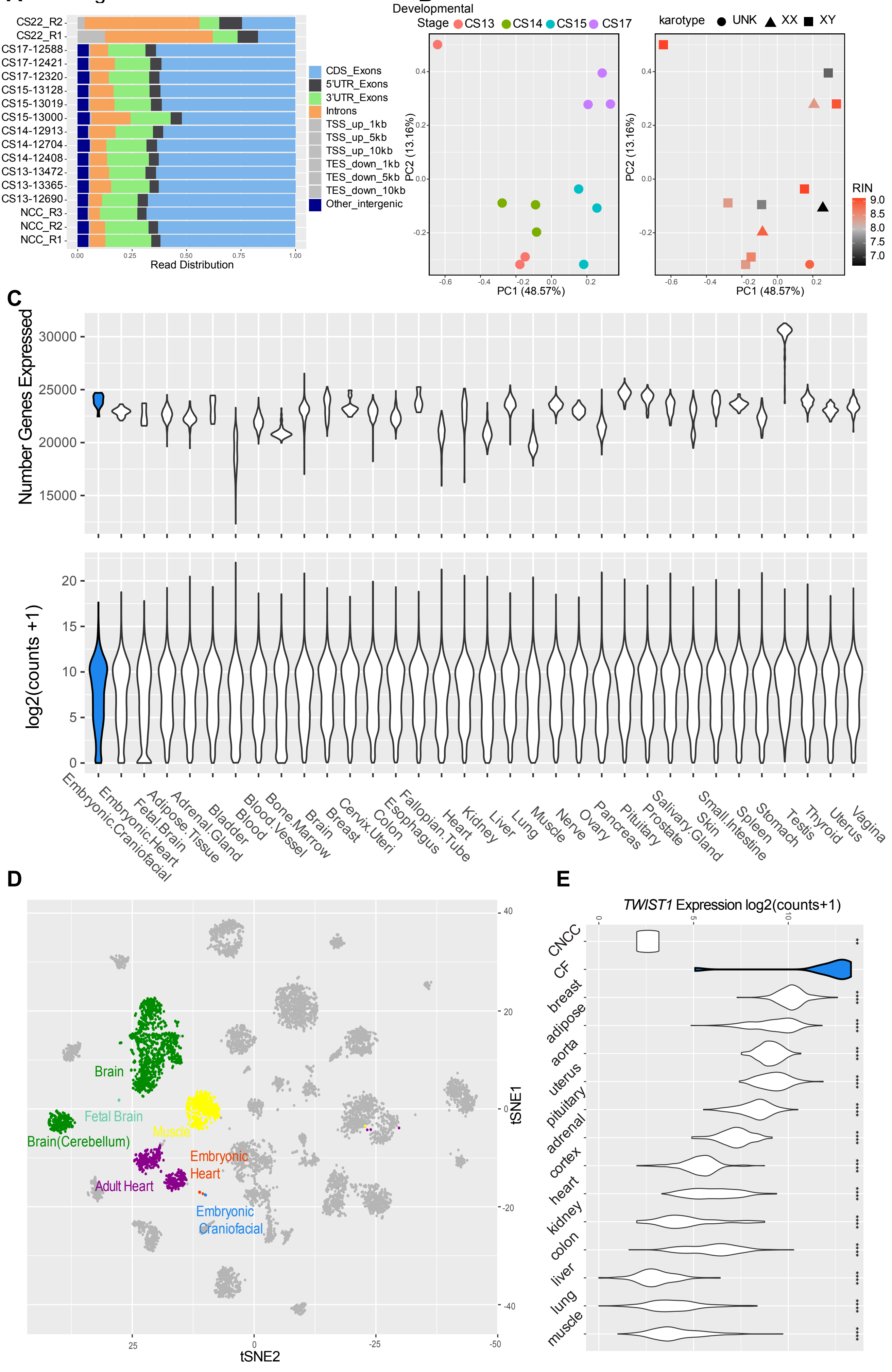
A
CS22_R2
CS22_R1
CS17-12588
CS17-12421
CS17-12320
CS15-13128
CS15-13019
CS15-13000
CS14-12913
CS14-12704
CS14-12408
CS13-13472
CS13-13365
CS13-12690
NCC_R3
NCC_R2
NCC_R1
Read Distribution
CDS_Exons
5'UTR_Exons
3'UTR_Exons
Introns
TSS_up_1kb
TSS_up_5kb
TSS_up_10kb
TES_down_1kb
TES_down_5kb
TES_down_10kb
Other_intergenic
B
Developmental Stage CS13 CS14 CS15 CS17
karotype UNK XX XY
PC2 (13.16%)
PC1 (48.57%)
RIN
C
Number Genes Expressed
log2(counts +1)
Embryonic.Craniofacial
Embryonic.Heart
Fetal.Brain
Adipose.Tissue
Adrenal.Gland
Bladder
Blood
Blood.Vessel
Bone.Marrow
Brain
Breast
Cervix.Uteri
Colon
Esophagus
Fallopian.Tube
Heart
Kidney
Liver
Lung
Muscle
Nerve
Ovary
Pancreas
Pituitary
Prostate
Salivary.Gland
Skin
Small.Intestine
Spleen
Stomach
Testis
Thyroid
Uterus
Vagina
D
Brain
Fetal Brain
Brain(Cerebellum)
Muscle
Embryonic Heart
Adult Heart
Embryonic Craniofacial
tSNE1
tSNE2
E
TWIST1 Expression log2(counts+1)
CNCC
CF
breast
adipose
aorta
uterus
pituitary
adrenal
cortex
heart
kidney
colon
liver
lung
muscle

A Figure S2

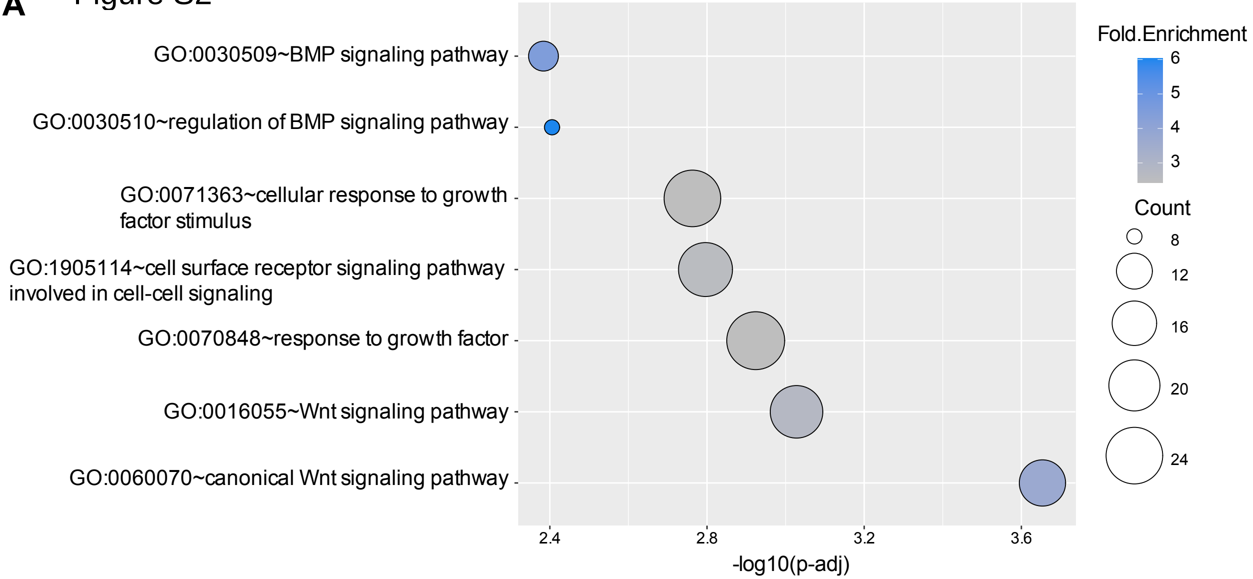

GO:0030509~BMP signaling pathway
GO:0030510~regulation of BMP signaling pathway
GO:0071363~cellular response to growth factor stimulus
GO:1905114~cell surface receptor signaling pathway involved in cell-cell signaling
GO:0070848~response to growth factor
GO:0016055~Wnt signaling pathway
GO:0060070~canonical Wnt signaling pathway
2.4
2.8
3.2
3.6
-log10(p-adj)
Fold.Enrichment
6
5
4
3
Count
8
12
16
20
24

### Figure S3

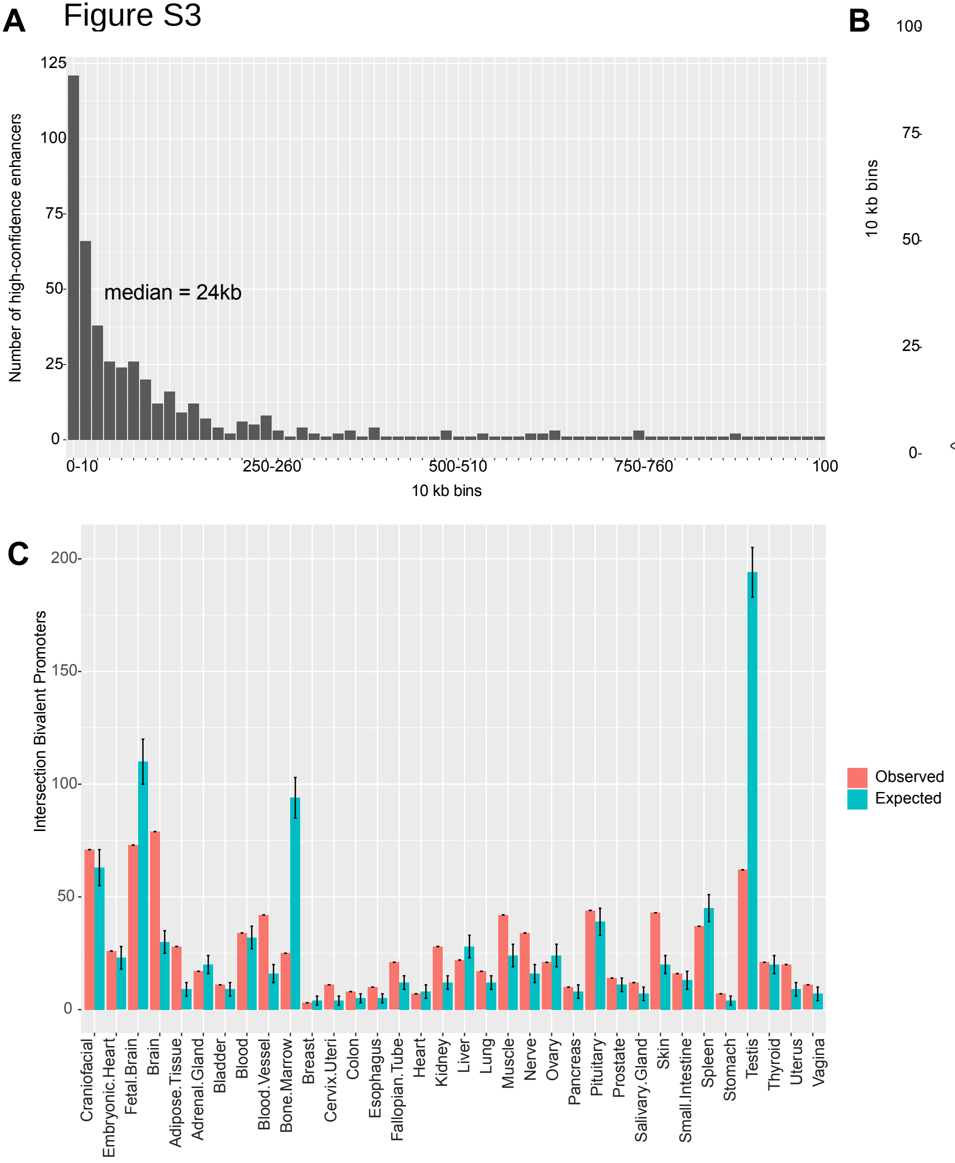

A
125
100
75
50
25
0
Number of high-confidence enhancers
median = 24kb
0-10
250-260
500-510
750-760
100
10 kb bins
B
100
75
50
25
0
10 kb bins
Gasperini et. al 2019
This study
C
200
150
100
50
0
Intersection Bivalent Promoters
Craniofacial
Embryonic.Heart
Fetal.Brain
Brain
Adipose.Tissue
Adrenal.Gland
Bladder
Blood
Blood.Vessel
Bone.Marrow
Breast
Cervix.Uteri
Colon
Esophagus
Fallopian.Tube
Heart
Kidney
Liver
Lung
Muscle
Nerve
Ovary
Pancreas
Pituitary
Prostate
Salivary.Gland
Skin
Small.Intestine
Spleen
Stomach
Testis
Thyroid
Uterus
Vagina
Observed
Expected

Figure S4

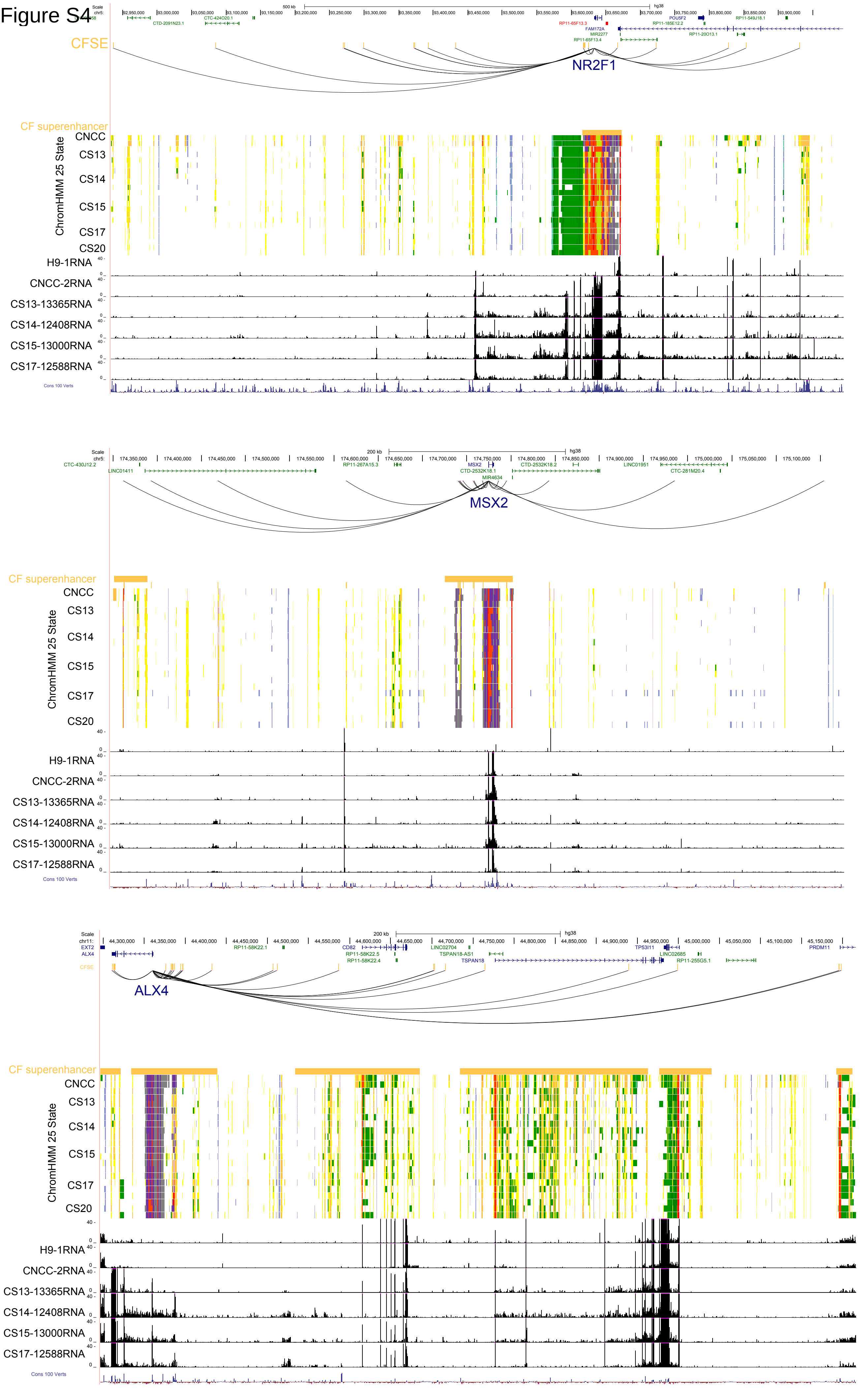

CFSE
NR2F1
CF superenhancer
ChromHMM 25 State
CNCC
CS13
CS14
CS15
CS17
CS20
H9-1RNA
CNCC-2RNA
CS13-13365RNA
CS14-12408RNA
CS15-13000RNA
CS17-12588RNA
Cons 100 Verts
MSX2
CF superenhancer
ChromHMM 25 State
CNCC
CS13
CS14
CS15
CS17
CS20
H9-1RNA
CNCC-2RNA
CS13-13365RNA
CS14-12408RNA
CS15-13000RNA
CS17-12588RNA
Cons 100 Verts
CFSE
ALX4
CF superenhancer
ChromHMM 25 State
CNCC
CS13
CS14
CS15
CS17
CS20
H9-1RNA
CNCC-2RNA
CS13-13365RNA
CS14-12408RNA
CS15-13000RNA
CS17-12588RNA
Cons 100 Verts

Figure S5

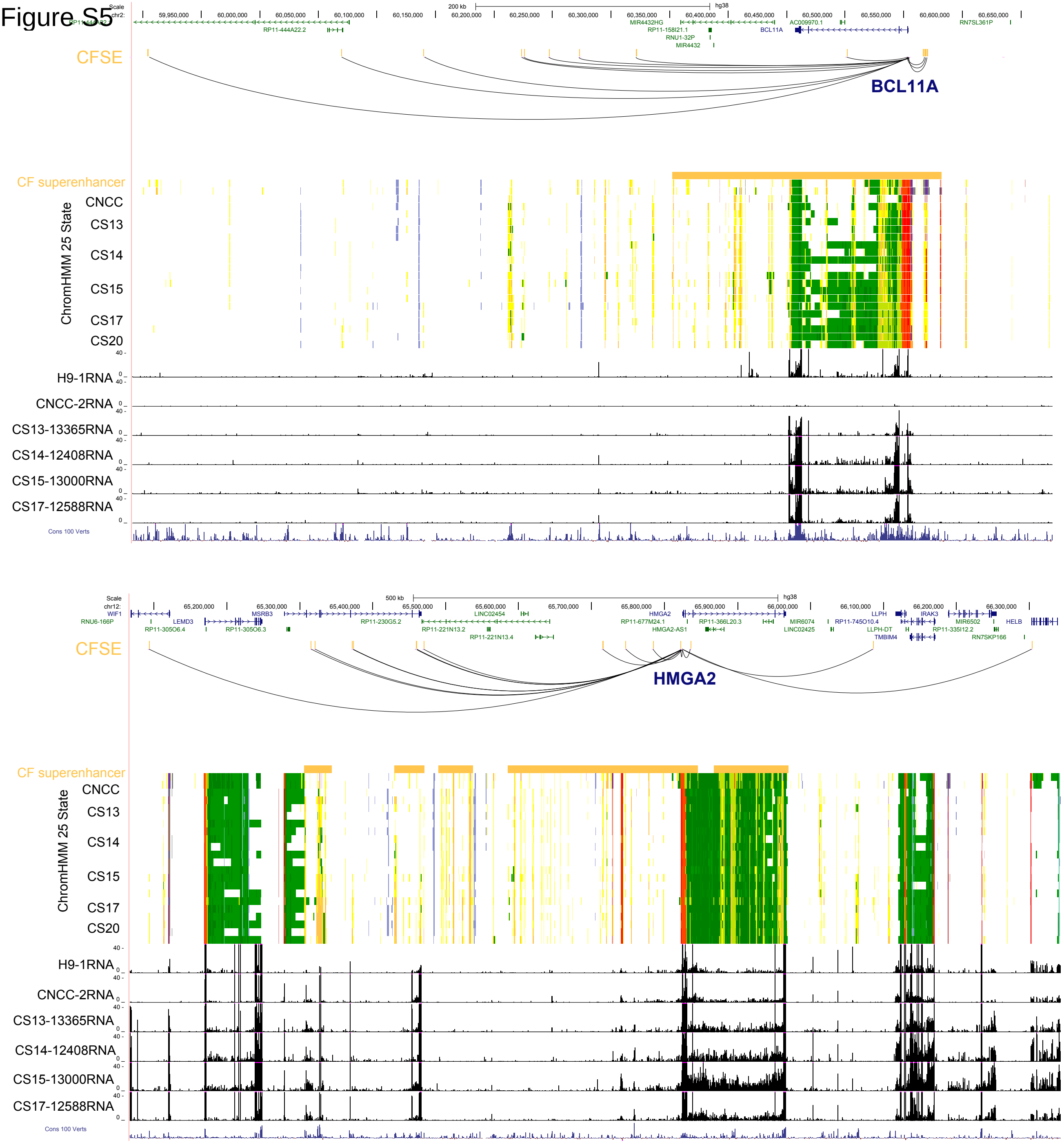
Scale
chr2:
200 kb
hg38
59,950,000
60,000,000
60,050,000
60,100,000
60,150,000
60,200,000
60,250,000
60,300,000
60,350,000
60,400,000
60,450,000
60,500,000
60,550,000
60,600,000
60,650,000
RP11-444A22.2
MIR4432HG
RP11-158I21.1
RNU1-32P
MIR4432
BCL11A
AC009970.1
RN7SL361P
CFSE
BCL11A
CF superenhancer
ChromHMM 25 State
CNCC
CS13
CS14
CS15
CS17
CS20
H9-1RNA
CNCC-2RNA
CS13-13365RNA
CS14-12408RNA
CS15-13000RNA
CS17-12588RNA
Cons 100 Verts
Scale
chr12:
500 kb
hg38
65,200,000
65,300,000
65,400,000
65,500,000
65,600,000
65,700,000
65,800,000
65,900,000
66,000,000
66,100,000
66,200,000
66,300,000
WIF1
RNU6-166P
LEMD3
MSRB3
RP11-305O6.4
RP11-305O6.3
RP11-230G5.2
LINC02454
RP11-221N13.2
RP11-221N13.4
HMGA2
RP11-677M24.1
HMGA2-AS1
RP11-366L20.3
MIR6074
LINC02425
LLPH
RP11-745O10.4
LLPH-DT
TMBIM4
IRAK3
MIR6502
RP11-335I12.2
RN7SKP166
HELB
CFSE
HMGA2
CF superenhancer
ChromHMM 25 State
CNCC
CS13
CS14
CS15
CS17
CS20
H9-1RNA
CNCC-2RNA
CS13-13365RNA
CS14-12408RNA
CS15-13000RNA
CS17-12588RNA
Cons 100 Verts

Figure S6

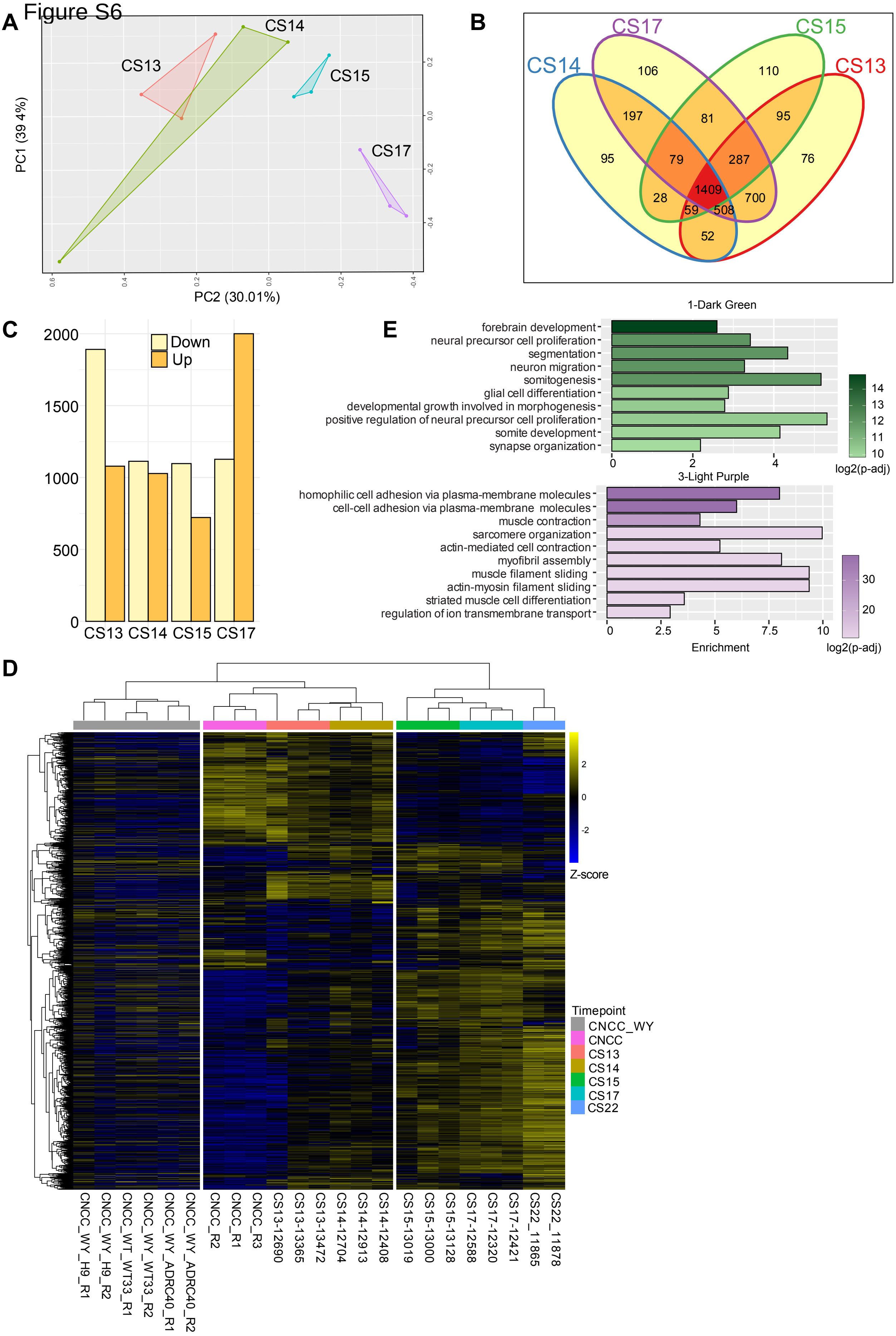
A
CS14
CS13
CS15
CS17
PC1 (39.4%)
PC2 (30.01%)
B
CS17
CS15
CS14
CS13
106
110
197
81
95
95
79
287
76
1409
28
700
59
508
52
C
Down
Up
2000
1500
1000
500
0
CS13 CS14 CS15 CS17
E
1-Dark Green
forebrain development
neural precursor cell proliferation
segmentation
neuron migration
somitogenesis
glial cell differentiation
developmental growth involved in morphogenesis
positive regulation of neural precursor cell proliferation
somite development
synapse organization
log2(p-adj)
3-Light Purple
homophilic cell adhesion via plasma-membrane molecules
cell-cell adhesion via plasma-membrane molecules
muscle contraction
sarcomere organization
actin-mediated cell contraction
myofibril assembly
muscle filament sliding
actin-myosin filament sliding
striated muscle cell differentiation
regulation of ion transmembrane transport
Enrichment
log2(p-adj)
D
Z-score
Timepoint
CNCC_WY
CNCC
CS13
CS14
CS15
CS17
CS22
CNCC_WY_H9_R1
CNCC_WY_H9_R2
CNCC_WT_WT33_R1
CNCC_WY_WT33_R2
CNCC_WY_ADRC40_R1
CNCC_WY_ADRC40_R2
CNCC_R2
CNCC_R1
CNCC_R3
CS13-12690
CS13-13365
CS13-13472
CS14-12704
CS14-12913
CS14-12408
CS15-13019
CS15-13000
CS15-13128
CS17-12588
CS17-12320
CS17-12421
CS22_11865
CS22_11878

### Figure S7

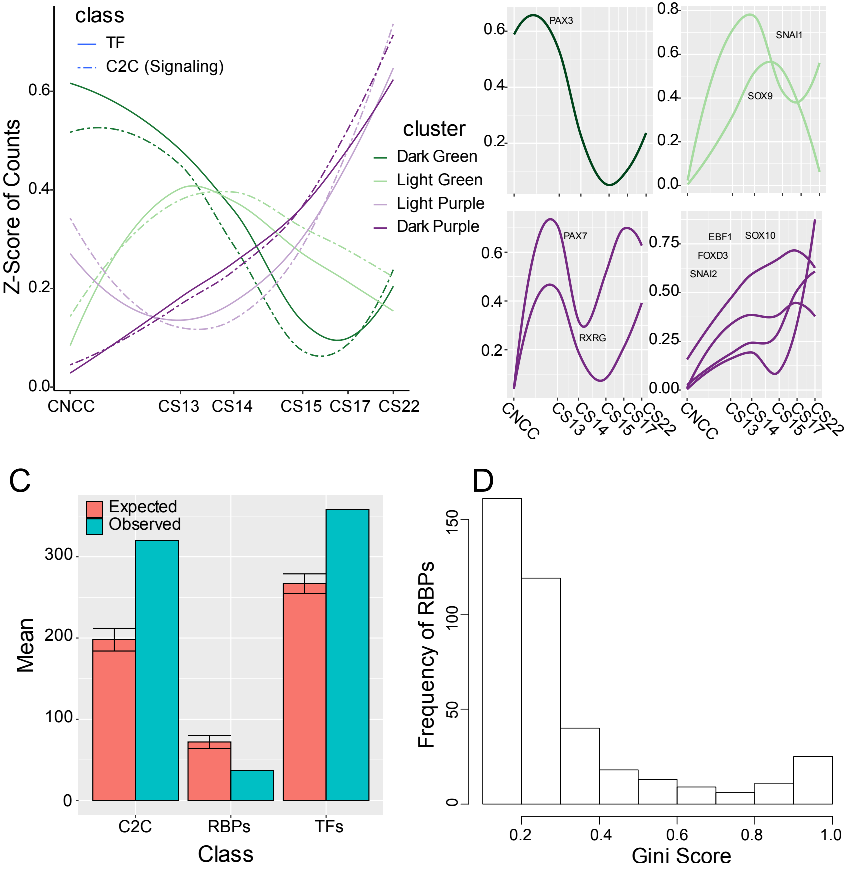
A
class
TF
C2C (Signaling)
cluster
Dark Green
Light Green
Light Purple
Dark Purple
Z-Score of Counts
0.0
0.2
0.4
0.6
CNCC
CS13
CS14
CS15
CS17
CS22
B
PAX3
SNAI1
SOX9
PAX7
RXRG
EBF1
SOX10
FOXD3
SNAI2
C
Expected
Observed
Mean
0
100
200
300
C2C
RBPs
TFs
Class
D
Frequency of RBPs
0
50
100
150
0.2
0.4
0.6
0.8
1.0
Gini Score

Figure S8

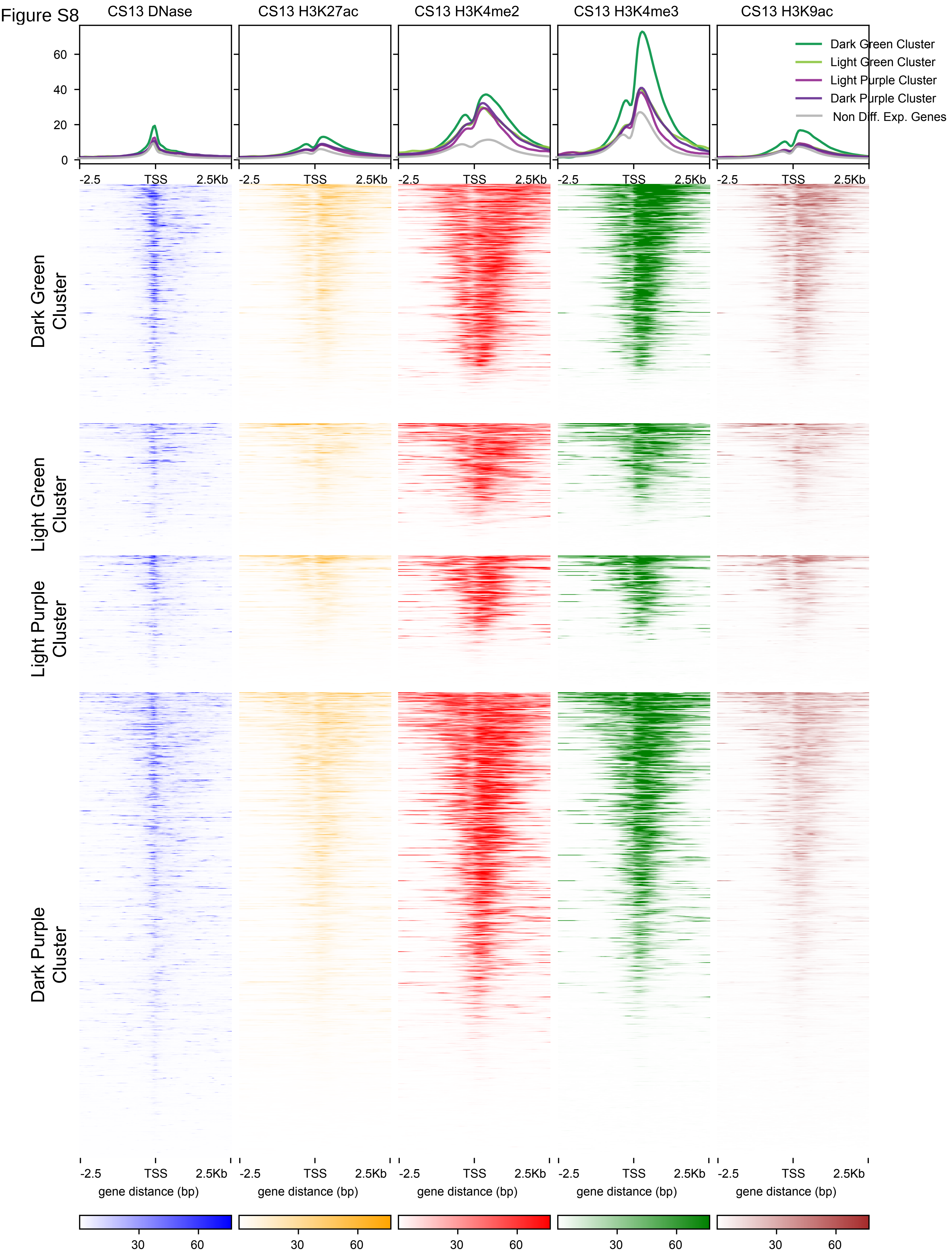
CS13 DNase
CS13 H3K27ac
CS13 H3K4me2
CS13 H3K4me3
CS13 H3K9ac
Dark Green Cluster
Light Green Cluster
Light Purple Cluster
Dark Purple Cluster
Non Diff. Exp. Genes
60
40
20
0
-2.5
TSS
2.5Kb
Dark Green Cluster
Light Green Cluster
Light Purple Cluster
Dark Purple Cluster
gene distance (bp)
30
60

### Figure S9

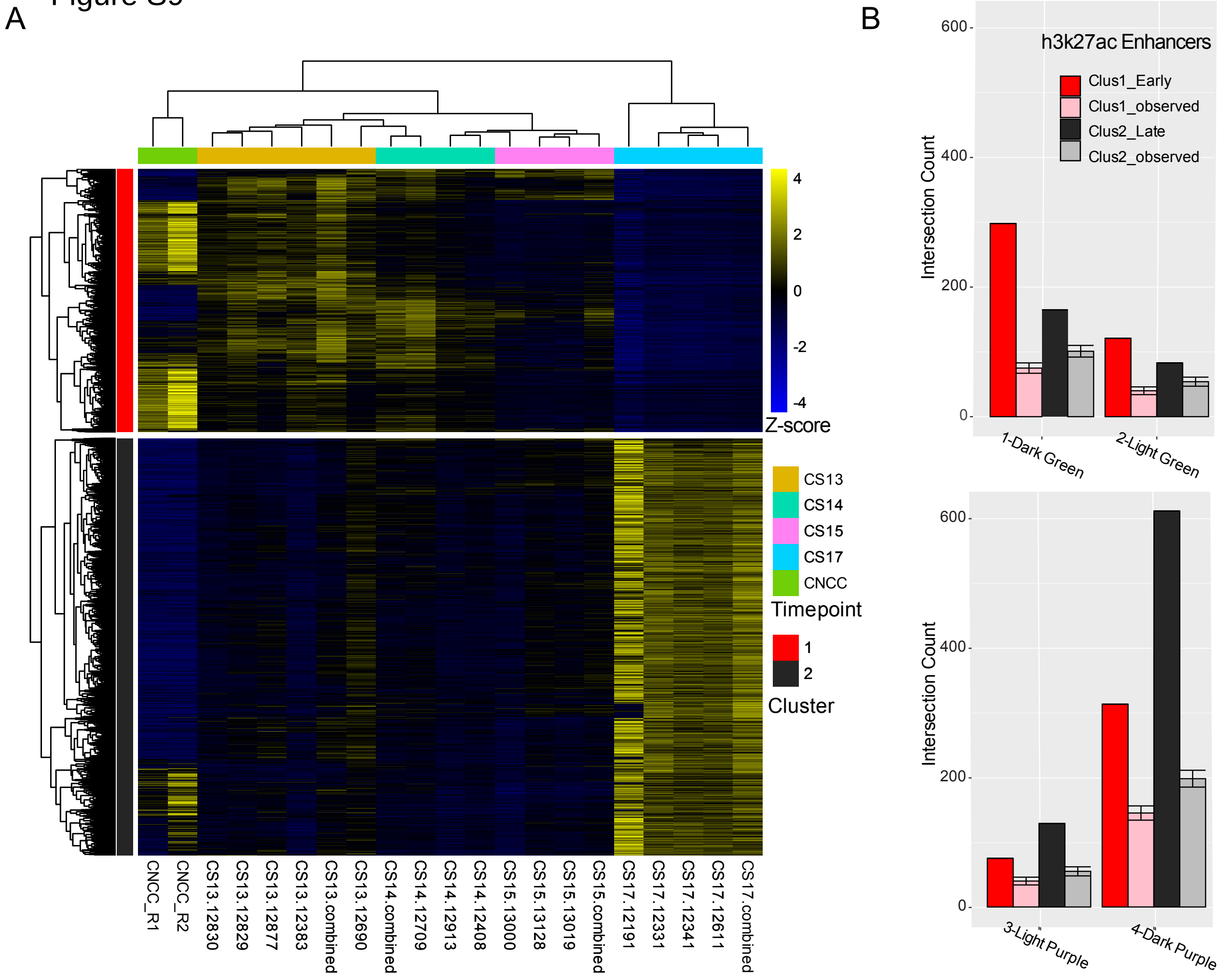

A
CNCC_R1
CNCC_R2
CS13.12830
CS13.12829
CS13.12877
CS13.12383
CS13.combined
CS13.12690
CS14.combined
CS14.12709
CS14.12913
CS14.12408
CS15.13000
CS15.13128
CS15.13019
CS15.combined
CS17.12191
CS17.12331
CS17.12341
CS17.12611
CS17.combined
4
2
0
-2
-4
Z-score
CS13
CS14
CS15
CS17
CNCC
Timepoint
1
2
Cluster
B
h3k27ac Enhancers
Clus1_Early
Clus1_observed
Clus2_Late
Clus2_observed
Intersection Count
600
400
200
0
1-Dark Green
2-Light Green
3-Light Purple
4-Dark Purple

Figure S10

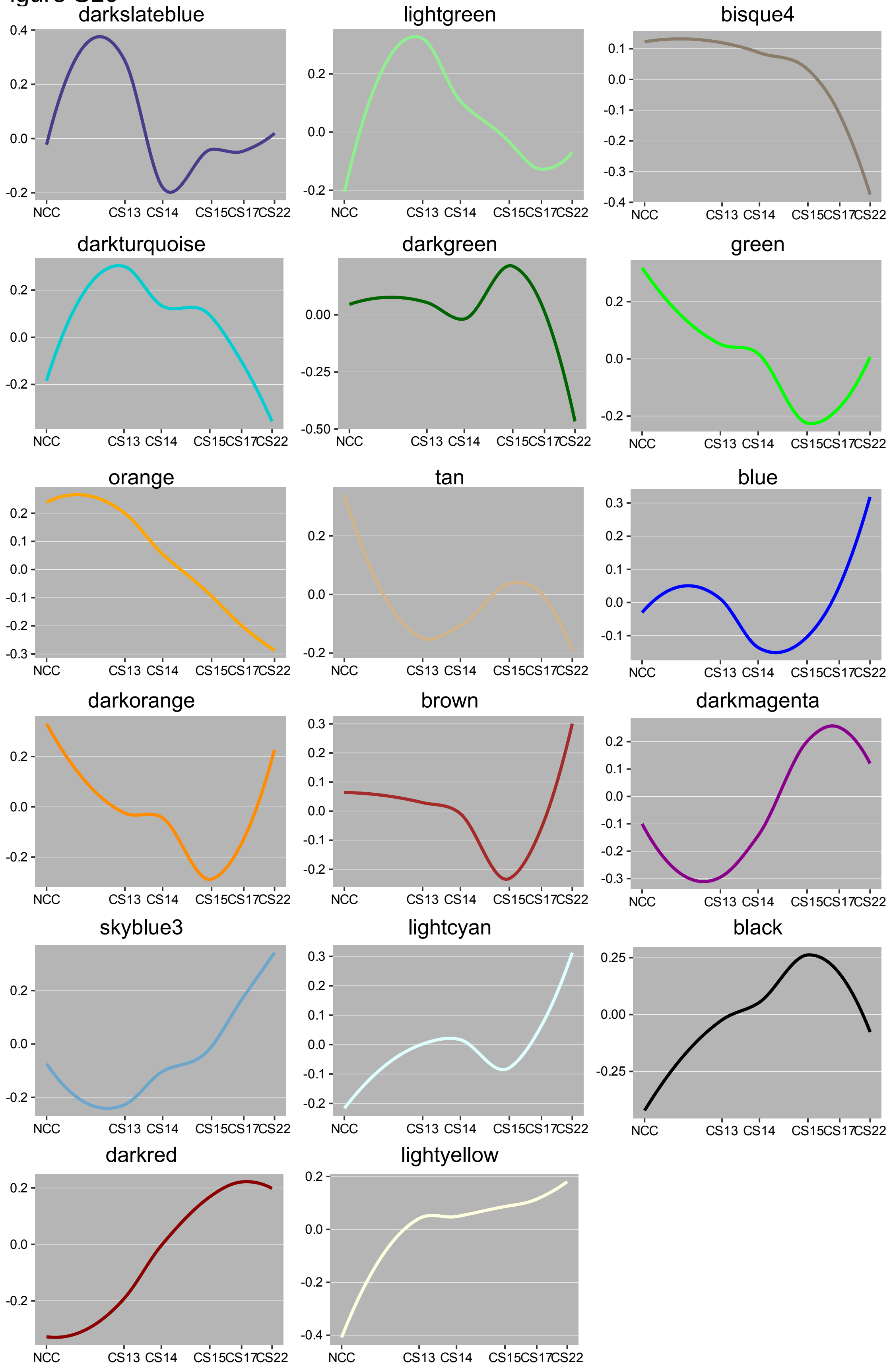
darkslateblue
lightgreen
bisque4
darkturquoise
darkgreen
green
orange
tan
blue
darkorange
brown
darkmagenta
skyblue3
lightcyan
black
darkred
lightyellow
NCC
CS13
CS14
CS15
CS17
CS22

Figure S11

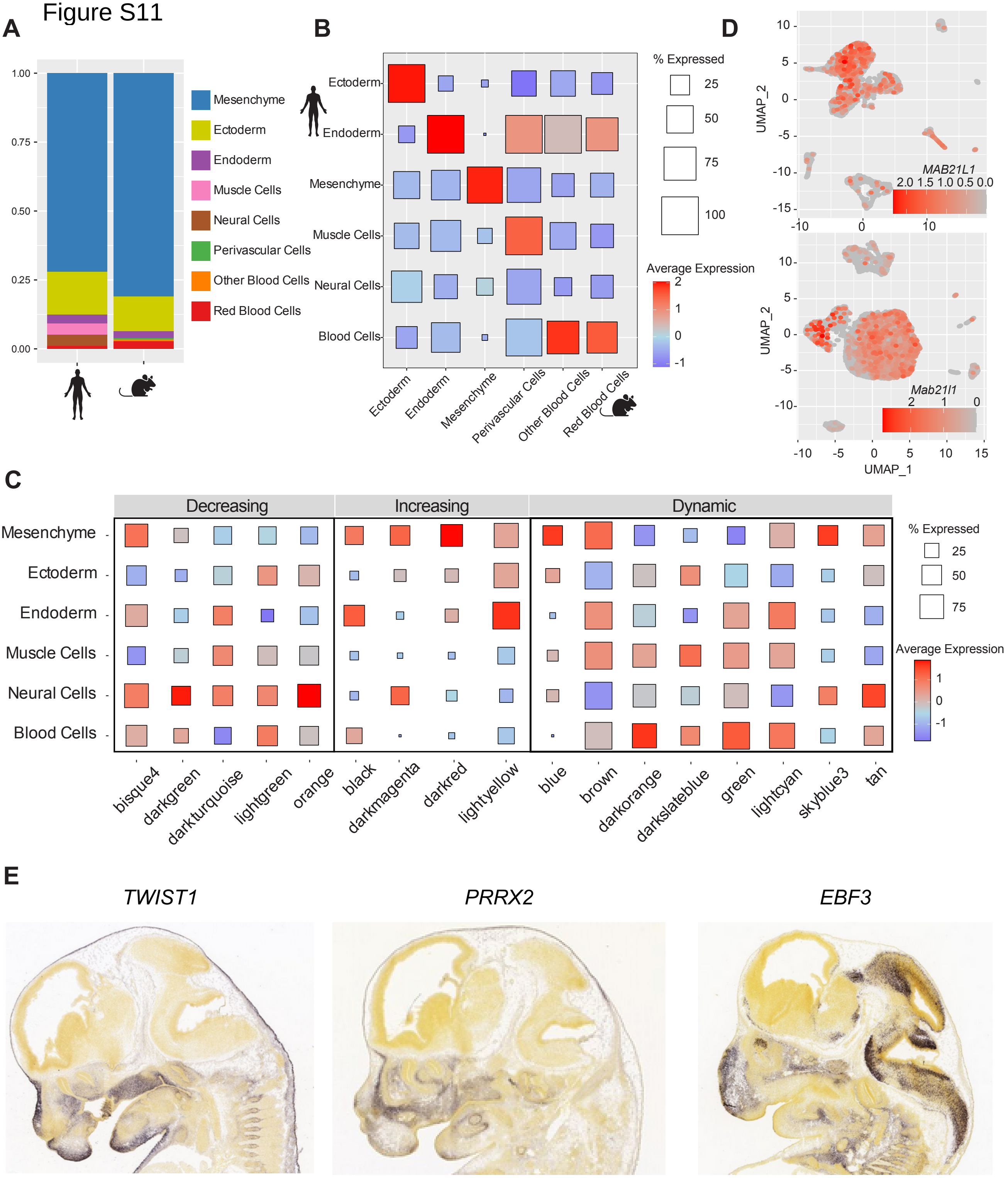

A
1.00
0.75
0.50
0.25
0.00
Mesenchyme
Ectoderm
Endoderm
Muscle Cells
Neural Cells
Perivascular Cells
Other Blood Cells
Red Blood Cells
B
Ectoderm
Endoderm
Mesenchyme
Muscle Cells
Neural Cells
Blood Cells
Ectoderm
Endoderm
Mesenchyme
Perivascular Cells
Other Blood Cells
Red Blood Cells
% Expressed
25
50
75
100
Average Expression
2
1
0
-1
D
MAB21L1
2.0 1.5 1.0 0.5 0.0
Mab21l1
2 1 0
UMAP_2
UMAP_1
C
Decreasing
Increasing
Dynamic
Mesenchyme
Ectoderm
Endoderm
Muscle Cells
Neural Cells
Blood Cells
bisque4
darkgreen
darkturquoise
lightgreen
orange
black
darkmagenta
darkred
lightyellow
blue
brown
darkorange
darkslateblue
green
lightcyan
skyblue3
tan
% Expressed
25
50
75
Average Expression
1
0
-1
E
TWIST1
PRRX2
EBF3
